# Circular Smad1-Encoded Polypeptide Regulates Myogenesis

**DOI:** 10.64898/2026.02.26.708385

**Authors:** Tanvi Sinha, Swarnava Dutta, Punit Prasad, Amaresh C. Panda

## Abstract

The majority of RNAs transcribed from the genome are non-coding RNAs (ncRNAs) that are involved in regulating the expression of protein-coding genes. However, a growing body of research highlights several novel microproteins encoded by unconventional ncRNAs such as long non-coding RNAs and circular RNAs (circRNAs) as important regulators of disease and development. Although several circRNAs have recently been reported to translate into functional peptides in diverse tissues, their roles in skeletal muscle remain largely unexplored. In this study, polyribosome-associated RNA sequencing and publicly available translatable circRNAs from the riboCIRC database were curated to discover potential protein-coding circRNAs in mouse C2C12 skeletal muscle cells. We validated a few circRNAs with high potential of translating into proteins in mouse C2C12 cells, including circular *Smad1* (*circSmad1*) that encodes a 194 amino acid peptide called circSmad1-194aa. Interestingly, silencing of *circSmad1* in C2C12 cells resulted in loss of myotube fusion and maturation. CircSmad1-194aa was found to contain the DNA-binding SMAD1-MH1 domain that localized into the nucleus during myogenesis. Moreover, CircSmad1-194aa associates with the BMP-responsive element (BRE) in the *Id1* promoter that is known to inhibit *Myod1*-driven myoblast differentiation. We propose that circSmad1-194aa promotes myogenesis by masking *Id1*-BRE from SMAD complex interaction, leading to suppression of ID1 expression and upregulation of MYOD1. Together, our findings identify circSmad1-194aa as a novel regulator of skeletal muscle differentiation and highlight the potential for the discovery of other functional circRNA-derived peptides in muscle pathophysiology.

**HIGHLIGHTS:**

- RNA-seq discovers hundreds of polysome-associated circRNAs in mouse C2C12 myoblasts
- Protein-coding circRNAs exhibit myogenesis-specific expression
- *CircSmad1* is abundant, methylated, and translated into a 194aa polypeptide
- *CircSmad1*-encoded polypeptide promotes myogenesis

**GRAPHICAL ABSTRACT:** 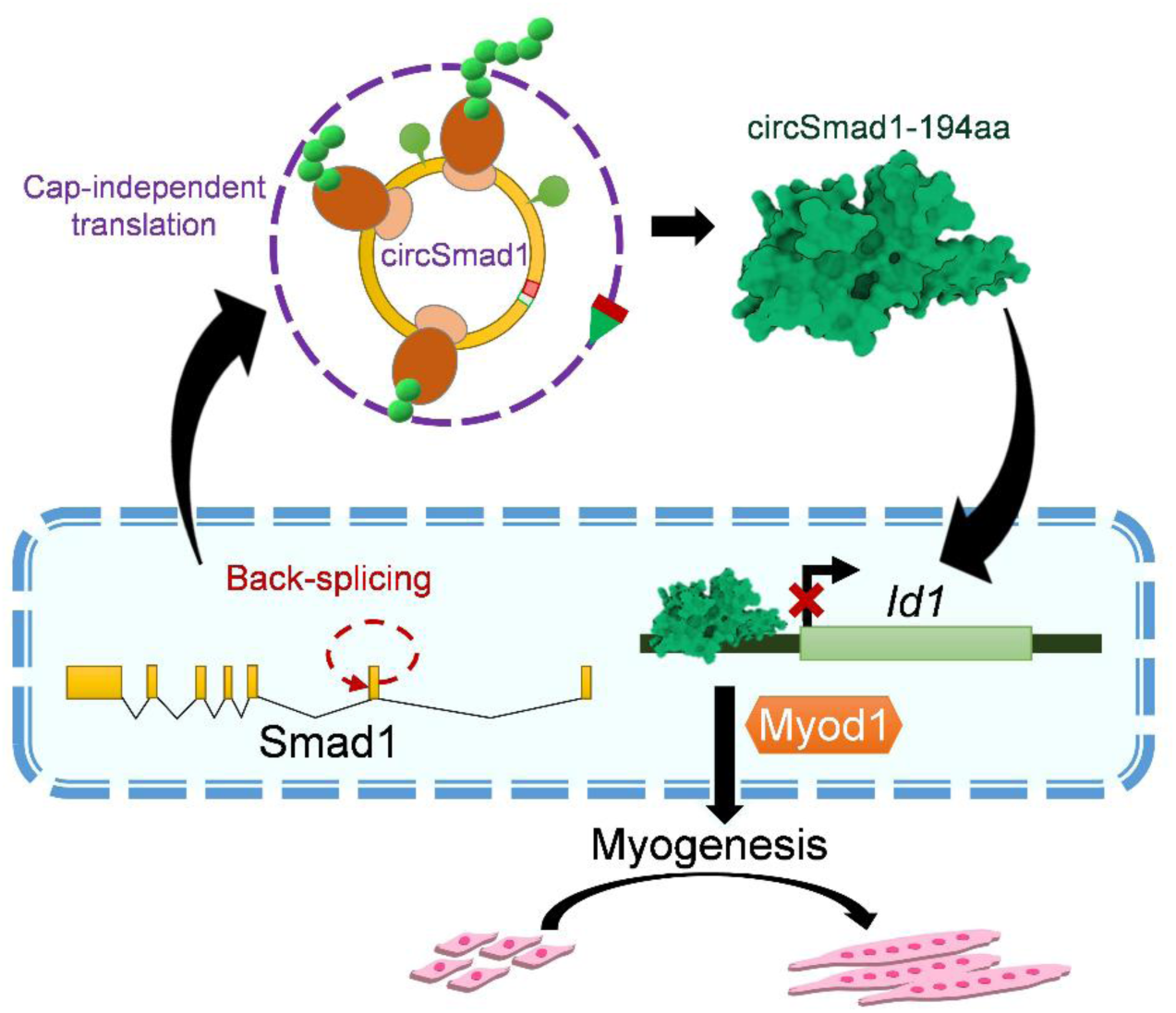

## INTRODUCTION

Skeletal muscle constitutes nearly 40% of total body mass and is essential for locomotion, metabolism, and systemic homeostasis [1, 2]. Its formation and regeneration rely on tightly regulated transcriptional and translational programs governed by myogenic regulatory factors (MRFs). The coordinated, stage-specific activity of MRFs ensures proper muscle development and adaptive responses to physiological demand and injury [3, 4]. Early MRFs, Myod1 and Myf5, specify muscle progenitors and initiate the myogenic program, whereas late MRFs, myogenin and MRF4, drive cell-cycle exit, myoblast fusion, and myofiber maturation [3, 4]. Although MRFs are well-studied for their function in myogenesis, recent advances in transcriptomics and proteomics have uncovered the roles of noncoding (nc)RNAs, including micro (mi)RNA, long noncoding (lnc)RNAs, and recently discovered circular (circ)RNAs [5–7].

CircRNAs represent a ubiquitously expressed family of conserved RNAs that is generated by head-to-tail splicing of pre-mRNAs [8]. CircRNAs were initially shown to function as non-coding regulatory RNAs by modulating transcription, and post-transcriptional events by interacting with miRNAs, mRNAs or RNA-binding proteins [9–11]. Although ncRNAs, including circRNAs, are mostly known for their gene regulatory role by interacting with other RNAs/proteins, emerging proteogenomic pipelines, combining ribosome profiling and custom mass spectrometry databases, are now uncovering thousands of previously unrecognised microproteins and non-canonical open reading frame (ncORF)-derived peptides [12–14]. Due to short length, rapid turnover, and technical challenges in detection, non-canonical proteins remain poorly characterized. [15].

Recent evidence suggests that microproteins play significant roles in myogenic differentiation, satellite cell activation, and metabolic regulation [16]. Although several studies continue to focus on the non-coding functions of circRNAs in myogenesis, many circRNAs have been shown to translate via cap-independent initiation through internal ribosome entry sites (IRESs) or N6-methyladenosine (m^6^A), yielding functional proteins [5, 9, 17–19]. The growing number of protein-coding circRNAs across the tree of life redefines their classification and highlights circRNAs as an underrepresented source of proteomic diversity [17]. The protein-coding potential of circRNAs in eukaryotes was first reported in 2017, showing translation of polysome-associated *circZNF609* producing a peptide that regulates myoblast proliferation [20]. Subsequently, recent studies discovered that *circFAM188B*-encoded circFAM188B-103aa, *circDdb1*-encoded circDdb1-867aa, and circular NEB-encoded 907-aa protein regulate muscle cell proliferation or differentiation in various organisms [21–23].

Despite technological advances, substantial gaps persist in our understanding of muscle-specific ncORFs, including circRNA-encoded peptides across development, regeneration, ageing, and disease. Given their enrichment during muscle development and inherent stability, circRNAs represent plausible substrates for endogenous translation, although the functional impact of circRNA-derived peptides in skeletal muscle remains largely unexplored. In this study, we compared circRNAs identified from polysome RNA sequencing with a list of putative protein-coding circRNAs available at the riboCIRC v1, finding hundreds of common translatable circRNAs in mouse C2C12 myoblasts [24]. We sought to validate a few putative protein-coding circRNAs through various biochemical assays, including their association with translating polyribosomes. Notably, we discovered circular *Smad1* (*circSmad1*), which harbours a junction-spanning ORF producing a 194 aa protein - circSmad1-194aa. Silencing of *circSmad1* in C2C12 cells resulted in loss of myotube fusion and maturation. Structural homology revealed the presence of a DNA-binding SMAD1-MH1 domain in circSmad1-194aa. In addition, we confirmed the nuclear localization of circSmad1-194aa, and its association with the BMP-responsive region in *Id1* (Inhibitor of DNA binding 1). Altogether, our investigation indicates that *circSmad1* regulates myogenesis by encoding circSmad1-194aa, which modulates *Id1* transcription as a dominant-negative moiety, thereby promoting muscle differentiation in C2C12 cells. The study demonstrates a novel regulatory node mediated by *circSmad1-*derived peptide. This research gives precedence for the discovery of similar circRNA-encoded peptides that regulate well-known myogenic pathways and identifies new avenues for clinical intervention.

## MATERIAL AND METHODS

### Cell culture and myogenic differentiation

Murine C2C12 myoblasts (ATCC) were maintained in high-glucose DMEM supplemented with 15% FBS and 1% penicillin-streptomycin at 37 °C in 5% CO_2_. Cells were differentiated at above 90% confluence in DMEM supplemented with 2% horse serum for four days or until myotubes were observed. Human skeletal muscle myoblasts (HSMM) were cultured in SkGM-2 BulletKit complete media according to the manufacturer’s recommendations (Lonza CC-2580-7) and differentiated as per the standard protocol.

### Giemsa-Jenner staining

Differentiating C2C12 cells were fixed in a 1:1 methanol-acetone mixture for 5 minutes, stained sequentially with Jenner working solution for 5 minutes and Giemsa for 10 minutes, rinsed in distilled water and air-dried. Myotube morphology was assessed in randomly selected microscopic fields according to the standard protocol [25].

### Polysome profiling and RNA preparation

Polysome fractionation of C2C12 cells was performed following a previously reported method with a few modifications [26, 27]. Briefly, 3-day differentiated C2C12 and HSMM myotubes were treated with cycloheximide (200 µg/ml, 10 minutes), followed by washing and lysis in polysome extraction buffer (PEB). Debris-free lysate was layered and fractionated into twelve parts (F1-F12) across a linear 10-50% sucrose density gradient by ultracentrifugation (Optima XPN-100, Beckman Coulter) with SW41Ti rotor. Polysome graph was monitored by the A_260_ absorbance pattern generated using the polysome flow cell machine (BioComp TRiAX). The polysome fractions (F6-F12) were pooled in RNA isolation reagent (RNA Biotech), and RNA was extracted using the phenol-chloroform precipitation method, followed by RNA sequencing [28].

### Library preparation and RNA sequencing

High-throughput RNA sequencing was performed to identify polysome-associated circRNAs in C2C12 cells. Input RNA integrity was determined using Bioanalyzer RNA 6000 Nano (Agilent). Ribosomal RNA-depleted libraries were prepared from C2C12 polysome-associated RNA using Collibri™ Stranded RNA Library Prep Kit (Thermo Fisher Scientific) according to the manufacturer’s protocol. Paired-end reads (100nt) were sequenced using the Illumina NovaSeq 6000 platform.

### Computational identification of circRNAs

Raw reads were quality assessed using FastQC (v2.7.11a) followed by adapter removal and trimming by fastp (v0.23.2) [29]. Clean reads were aligned to the mouse genome (GRCm39 vM36) using STAR (v2.7.11a) with --chimSegmentMin 10 parameter [30]. CircRNAs were identified using the CIRCexplorer2 (v2.3.8) pipeline with default parameters [31]. Spliced and genomic sequences of circRNAs were extracted using the BEDTools (v2.31.1) *getfasta* command-line **(Supplementary Table S1)** [32]. CircRNAs common between myoblast and myotube polysome RNA-seq libraries, with supporting translation evidence from riboCIRC v1 (accessed on 10^th^ March 2021), were selected for downstream analyses **(Supplementary Table S2)**.

### RT-qPCR and Sanger sequencing of polysome-associated circRNAs

Total RNA was extracted using the RNA isolation reagent (RNA Biotech) method, and reverse-transcribed with random hexamer primers using Maxima RT (Thermo Fisher Scientific) as per the manufacturer’s protocol. Divergent primers spanning the back-spliced junction were used to validate individual circRNAs by SYBR Green qPCR (Thermo Fisher Scientific). Divergent PCR products with or without RT were resolved on a SYBR Gold stained 2% agarose gel, followed by purification of amplified DNA and Sanger sequencing to confirm the co-transcriptional back-splicing of circRNAs in C2C12 cells. Relative expression was calculated using the delta Ct method and normalised to *Gapdh* or *18S* rRNA housekeeping control **(Supplementary Table S3)** [33]. Further, the distribution of control mRNA, lncRNA and target circRNAs across the monosome and polysome fractions of C2C12 myotubes was determined by RT-qPCR. Polysome occupancy of circRNA was calculated as the percentage of circRNA present in polysome fractions relative to the total input. All experiments were performed with at least three independent biological replicates.

### RNase R resistance and Actinomycin D chase assay

Total RNA was treated with RNase R (4U/µg RNA) for 30 minutes at 37 °C, followed by RT-qPCR to confirm circular topology or stability of circRNAs, following a previously reported method [34]. To assess circRNA stability, an actinomycin D (ActD) chase assay was performed as described previously, with a few changes [35]. Briefly, C2C12 myoblasts were treated with ActD (5 µg/mL) to inhibit *de novo* transcription, and total RNA was isolated at different time points (0h, 0.5h, 1h, 2h, 4h, and 8h). Target circRNAs and mRNA levels were measured by RT-qPCR and normalised to basal (0h) to calculate RNA decay rates. All experiments were performed with at least three independent biological replicates.

### Species conservation and structure of *circSmad1*

Conservation of *circSmad1* was assessed by multiple sequence alignment with cross-species homologs of *circSmad1* mentioned in CIRCpedia v3 using Clustal Omega (https://www.ebi.ac.uk/jdispatcher/msa/clustalo), and a phylogenetic tree was generated using iTOL [36–38]. Secondary structure of *circSmad1* was predicted using the RNAfold web server with circular structure-specific parameters [39]. N6-methyladenosine (m^6^A) modification sites in the *circSmad1* mature sequence were predicted using the SRAMP web tool, and supporting MeRIP-seq data were obtained from the m6A2Circ repository [40, 41].

### Structural homology of *circSmad1*-encoded peptide

The 3x concatenated full-length sequence of *circSmad1* was used to predict junction-spanning circular ORF (cORF) using the NCBI ORFfinder tool [42]. The 3D structure of circSmad1-194aa was predicted using AlphaFold2 Colab with default parameters [43]. Sequence similarity between the circSmad1-194aa and the host SMAD1 was predicted using BLASTp [44]. The presence of functional domains in circSmad1-194aa was predicted using InterProScan webserver (accessed on 3^rd^ December 2025) [45]. Structure alignment of circSmad1-194aa with the publicly available SMAD1-MH1 structure (pdb_00003kmp) was performed using the RCSB PDB webserver (https://www.rcsb.org/alignment) [46, 47]. The circSmad1-194aa sequence was compared with orthologous circSmad1-cORFs across mammals using MUSCLE (https://www.ebi.ac.uk/jdispatcher/msa/muscle) to assess species conservation [38].

### MS/MS detection of *circSmad1*-encoded peptide

CircRNAs identified from C2C12 polysome RNA-seq were matched with a list of putative protein-coding mouse circRNAs downloaded from riboCIRC v1 (accessed on 10^th^ March 2021) [24]. The publicly available proteome dataset (PXD023256) listed as supporting evidence for *circSmad1* translation in riboCIRC v1 was retrieved from the PRIDE database, and analyzed using MaxQuant software (v 2.6.2.0) to detect the junction-spanning peptide of *circSmad1* [48, 49].

### m⁶A RNA immunoprecipitation

Total RNA extracted from C2C12 myotubes was subjected to m⁶A RNA immunoprecipitation (MeRIP) as described previously [50]. Initially, 5 µg each of target anti-m⁶A (Abcam) and control anti-IgG (CST) antibodies were allowed to hybridise with Dynabeads Protein G-coated magnetic beads (Thermo Fisher Scientific) for 2 hours at room temperature, followed by incubation with 15 µg of total RNA at 4°C for 2 h. Immunoprecipitated and input RNA were analyzed by junction-specific RT-qPCR assay. The relative enrichment of *circSmad1* in the m⁶A pulldown sample was calculated relative to the IgG control. *CircZfp609* was used as a positive circRNA control for the MeRIP assay [20].

### CircSmad1 silencing

Antisense oligo Gapmer (50nM) specific to the back-spliced junction of *circSmad1* was transfected into C2C12 myoblasts at 90% confluence using Lipofectamine RNAiMAX reagent (Thermo Fisher Scientific), following the manufacturer’s protocol. Similarly, control cells were transfected with a non-targeting Gapmer. Transfected myoblasts were induced to differentiate into myotubes for 96 h, and total RNA was isolated using the RNA isolation reagent (RNA Biotech) method followed by RT-qPCR assay. Protein lysates of transfected myotubes were prepared in 1X RIPA buffer and assessed by immunoblotting with antibodies against muscle-specific markers, including MYOD1 (ABclonal), MYOG (Thermo Fisher Scientific), and MYH3 (SCBT). All experiments were performed in at least three biological replicates.

### CircSmad1-194aa overexpression

Mammalian expression vector pCMV6, containing the predicted 194aa open reading frame of *circSmad1* fused to a C-terminal 3×FLAG epitope, was used to overexpress circSmad1-194aa in skeletal muscle cells. C2C12 myoblasts were transfected with pCMV6-circSmad1 or pcDNA3 empty plasmid using Lipofectamine 3000 (Thermo Fisher Scientific) as per the manufacturer’s instructions. Myoblasts were harvested two days after transfection, followed by detection of FLAG-tagged circSmad1-194aa using immunoblotting with anti-FLAG (CST) antibody.

### Immunofluorescence

Immunofluorescence (IF) analysis was performed in C2C12 cells transfected with pCMV6-circSmad1, following a previous protocol with a few modifications [51]. C2C12 myoblasts 48 h post-transfection or after differentiation into 4-day myotubes were fixed, permeabilised, and blocked with 3% BSA. FLAG-tagged circSmad1-194aa was detected by indirect IF staining with anti-FLAG (CST) primary antibody followed by Alexa Fluor 488-conjugated secondary antibody. Myocyte enhancer factor-2 (MEF2C), a myogenic marker, was detected in myotubes incubated with anti-MEF2C primary antibody (Thermo Fisher Scientific) at 4 °C overnight, followed by staining with Alexa Fluor 594-conjugated secondary antibody. Nuclei were counterstained with DAPI in mounting media (Thermo Fisher Scientific), and images were acquired using confocal microscopy (Leica SP8 STED).

### Subcellular fractionation

Cytoplasmic fraction was prepared from C2C12 cells by resuspending the cell pellet in a hypotonic cytoplasmic lysis buffer (CLB), followed by RNA isolation with RNA isolation reagent (RNA Biotech), or used directly for the detection of cytosolic proteins by immunoblotting with suitable antibodies. The nuclear pellet was washed twice in CLB to remove residue and dissolved in RNA isolation reagent (RNA Biotech) to isolate nuclear RNA. Or else the nuclear pellet was incubated in 1X RIPA buffer to extract the soluble nuclear fraction (SNF), and the undissolved chromatin pellet was sonicated in 1X RIPA buffer to extract the insoluble nuclear fraction (INF). Subcellular fractionation was verified by differential enrichment of *Malat1* (nuclear) and *H19* (cytoplasmic) transcripts in RT-qPCR assay. Immunoblotting with anti-GAPDH (CST) (cytoplasmic) and anti-Lamin A/C (CST) (nuclear) antibodies was used to confirm the subcellular fractionation of C2C12 cells at the protein level.

### Cycloheximide chase assay

To determine the stability of circSmad1-194aa, transfected C2C12 cells were treated with cycloheximide (CHX) (50 µg/ml) and harvested at defined time intervals (0h, 2h, 4h, and 6h) as described previously [52]. The relative abundance of FLAG-tagged circSmad1-194aa was quantified by immunoblotting with anti-FLAG antibody (CST). After CHX treatment, the protein level at each time interval was measured relative to the basal level (0h).

### Chromatin immunoprecipitation

Chromatin immunoprecipitation (ChIP) was performed to examine in vivo DNA-binding activity of the circSmad1-194aa in C2C12 myoblasts, as described previously [53]. Briefly, two days after transfection, cells were cross-linked with formaldehyde, which was followed by nuclear isolation. Chromatin was sheared by sonication to an average fragment size of 200–500 bp. The cleared chromatin was incubated overnight with anti-FLAG antibody or control anti-IgG bound to Protein A/G magnetic beads (Thermo Fisher Scientific), and an aliquot was saved as input. After three washings, the bound DNA-protein complex was eluted, treated with Proteinase K, and the cross-links were reversed. DNA was purified and quantified by SYBR Green qPCR (Thermo Fisher Scientific) with *Id1*-BRE primers [54]. The *Id1*-BRE Ct values were normalised to input, and enrichment was estimated relative to IgG. The *18S* rRNA locus was used as a negative control.

### Biotinylated dsDNA-protein immunoprecipitation

A streptavidin-biotin dsDNA immunoprecipitation assay was conducted to evaluate the DNA-binding specificity of circSmad1-194aa in C2C12 cells. Myoblasts were lysed, 48 h post-transfection, under non-denaturing conditions, following a previous method with a few changes [55]. Both cytoplasmic and nuclear extracts were combined, and an aliquot was reserved as input [56]. A biotinylated dsDNA probe corresponding to the 29bp *Id1*-BRE region was generated by annealing complementary oligonucleotides through heating at 95°C, followed by gradual cooling to room temperature in annealing buffer [54]. A non-targeting biotinylated dsDNA probe was used as a negative control. An equal amount of cell lysate was incubated with *Id1*-BRE and control dsDNA probes, and DNA-protein complexes were captured using streptavidin-coated magnetic beads (Thermo Fisher Scientific). After extensive washing, bound proteins were eluted by boiling in 1X SDS loading buffer, and the circSmad1-FLAG was detected by immunoblotting with anti-FLAG antibody (CST). All experiments were performed with at least three independent biological replicates.

### Statistical analysis

All experiments were performed with at least three independent biological replicates and are mentioned in the figure legends. Statistical analyses were performed using GraphPad Prism or Microsoft Excel. Significance was determined using the two-tailed Student’s t-tests, with p-value <0.05 considered significant unless otherwise stated.

## RESULTS

### Thousands of circRNAs are associated with translating polyribosomes in C2C12 cells

To study the role of protein-coding circRNAs in myogenesis, we used the proliferating myoblasts (MB) and differentiated myotubes (MT) of cultured mouse C2C12 cells (**Figure 1A**). The expression of well-known myogenic markers, including Myod1, Myogenin (Myog), and myosin heavy chain (Myhc), was determined by both RT-qPCR and western blotting to confirm muscle differentiation in C2C12 cells (**Figure 1B,C**). To assess the landscape of polyribosome-associated circRNAs in C2C12 cells, we fractionated cycloheximide-treated C2C12 myoblasts and myotubes (**Figure 1D,E**). Total RNA was isolated from a pool of polysome fractions (F6-F12) and subjected to RNA sequencing to identify polysome-bound RNAs in C2C12 cells. Interestingly, analysis of RNA-seq data using the CIRCexplorer2 pipeline identified more than 3,700 circRNAs in differentiating C2C12 polysome samples (**Supplementary Table S1, Figure 1F**). We compared the circRNAs identified in the MB and MT polysome datasets with circRNAs reported in the riboCIRC v1 database, yielding a list of 76 common putative translatable circRNAs in mouse skeletal muscle (**Supplementary Table S2, Figure 1G**). Almost half of the circRNA population was composed of five or fewer exons or introns with a spliced length less than a thousand nucleotides (**Figure 1H, I**). Nearly eighty per cent of the polysome-bound circRNAs originated from exonic regions of the genome, suggesting possible open-reading frame overlap with the host gene (**Figure 1J**).

**Figure 1:**
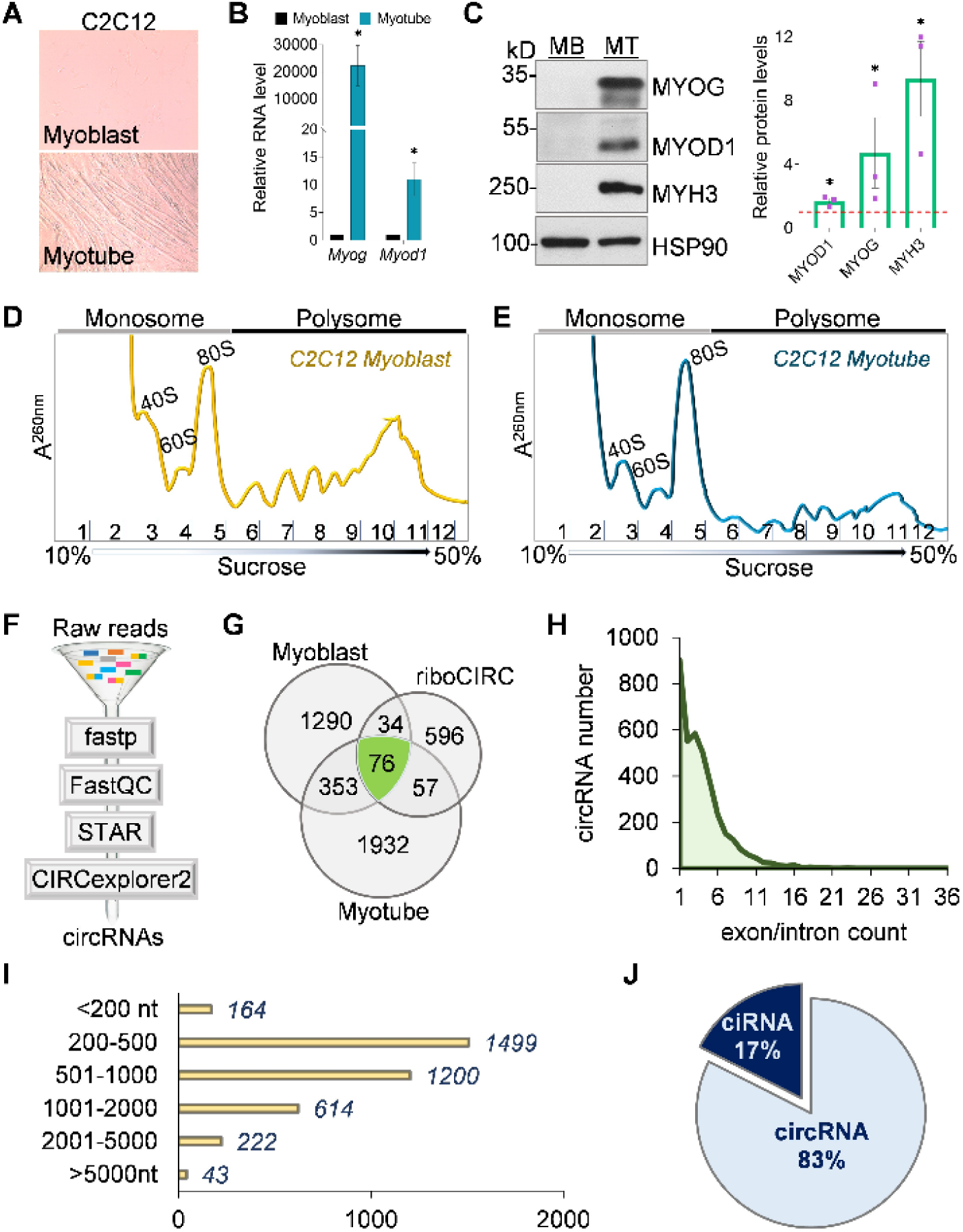
Identification of polysome-associated circRNAs in C2C12 cells. **A.** Representative images of myoblast and myotube in C2C12 cells. **B.** Bar chart showing levels of myogenic markers *Myog* and *Myod1* in differentiating C2C12 cells, normalised to *Gapdh* mRNA by RT-qPCR. **C.** Representative immunoblot detecting bands of MYOG, MYOD1 and MYH3 in C2C12 MB, MT (left), and their relative levels in myotubes normalised to HSP90 loading control shown in the densitometry bar plot (right). **D-E.** Representative polysome profiling graphs of RNA in different fractions of C2C12 myoblasts (left) and myotubes (right), distinguishing the monosome (F1-F6) and polysome (F7-F12) fractions. **F.** Schematic diagram showing bioinformatic workflow used for the identification of circRNAs from RNA-seq read input. **G.** Venn diagram showing the number of common and exclusive circRNAs between myoblast, myotube and riboCIRC v1. **H.** Area plot showing the distribution of circRNA population depending on the number of exons or introns. **I.** Bar plot showing the grouping of circRNAs based on spliced length range. **J.** Pie chart showing the percentage of the identified circRNAs that originate from exonic (circRNAs) or intronic (ciRNAs) regions of the genome. Data displayed in panels B and C represent the mean ± SEM of at least 3 biological replicates, and * indicates significance with a p-value <0.05.

### Polysome-bound circRNAs are differentially expressed during muscle cell differentiation

Polysome-associated circRNAs with high abundance in C2C12 cells that are already reported in the riboCIRC v1 database were filtered based on size (<2kb), exon count (<4), and multi-omics evidence (RPF, cORF, MS, m^6^A) available at riboCIRC to identify high-confidence protein-coding circRNAs (**Supplementary Table S2**) [24]. From the stringent screening, we selected five potential protein-coding circRNAs (Figure 2A). We validated the expression of the selected circRNAs in C2C12 cells using RT and no-RT PCR with divergent primers (**Supplementary Table S3**). Candidate circRNAs were detected as specific bands in RT-PCR, which were distinctly absent in the no-RT control, confirming the non-genomic or transcriptomic origin of the circRNAs in C2C12 cells (Figure 2B). Moreover, we purified and sequenced the RT-PCR products, confirming the occurrence of a back-splicing event unique to each circRNA in mouse skeletal muscle cells (Figure 2C). To check the circular conformation of circRNAs in C2C12 cells, we performed the RNase R resistance assay. We confirmed that the candidate circRNAs were not digested, while the linear *Gapdh* and *Actb* mRNAs were depleted due to active exoribonuclease degradation by RNase R compared to mock (Figure 2D). Myogenesis involves well-regulated changes in the expression of various coding and non-coding genes, including circRNAs, thereby maintaining muscle homeostasis [5, 57]. To assess the role of potential protein-coding circRNAs in myogenesis, we investigated the expression of circRNAs in C2C12 myoblasts and myotubes using RT-qPCR. We found that two polysome-associated circRNAs - *circSmad1* and *circH6pd* were significantly upregulated upon differentiation of C2C12 cells, suggesting a possible function of polysome-bound circRNAs in mouse myoblast differentiation (Figure 2E). Further, we investigated the distribution of *circSmad1* and *circH6pd* between the monosome and polysome fractions of C2C12 myotubes using RT-qPCR. *Gapdh* and *Myog* mRNAs were used as positive controls. Both *circSmad1* and *circH6pd* showed relative levels above 5% in polysome, suggesting the protein-coding potential of polysome-loaded circRNAs, consistent with the RNA-seq data (Figure 2F).

**Figure 2:**
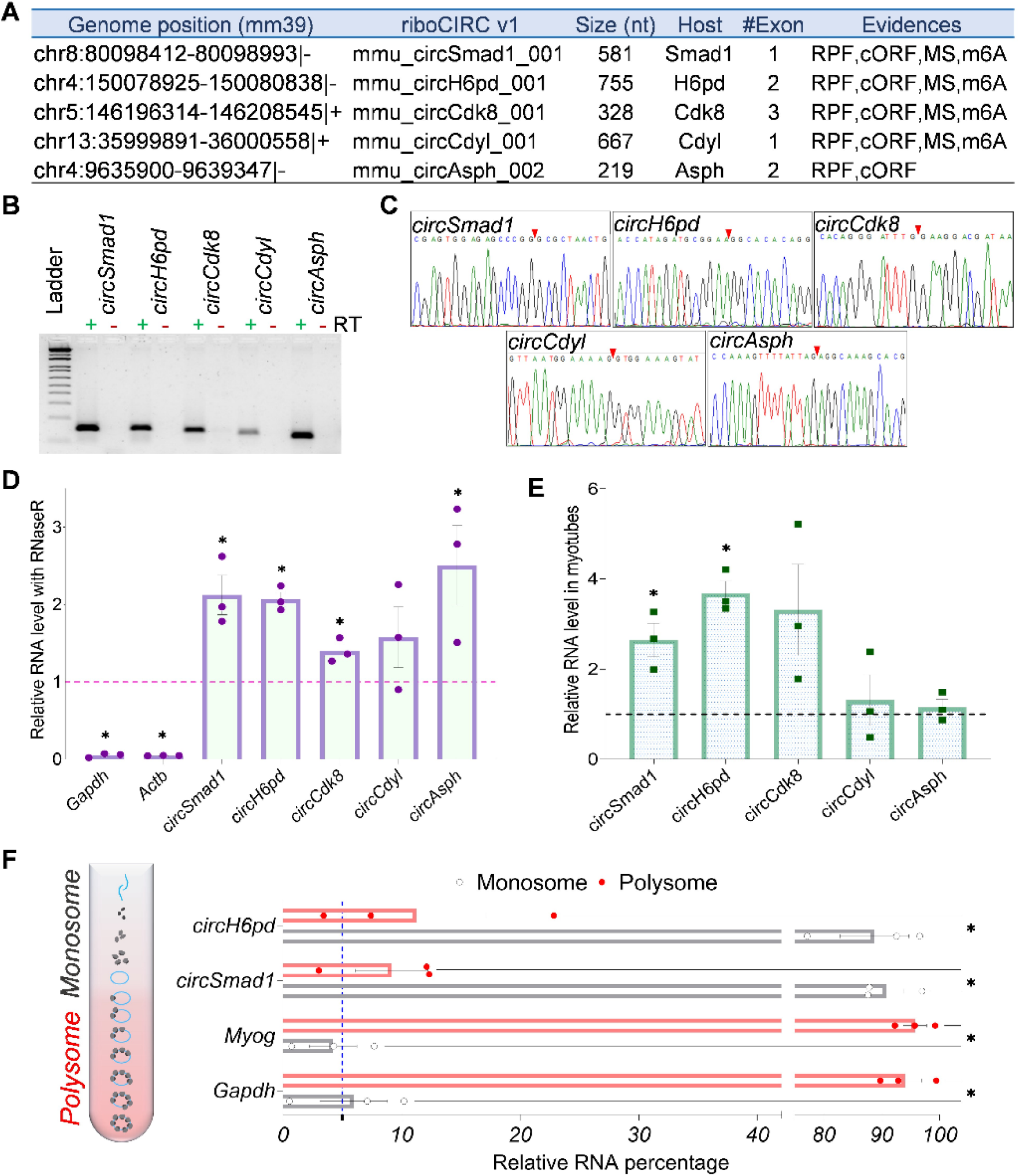
Validation of polysome-associated circRNAs in C2C12 cells. **A.** Table enlisting five C2C12 polysome-bound circRNAs and corresponding coding evidence retrieved from the riboCIRC v1 database. **B.** Agarose gel image showing divergent PCR bands present in RT (+) and absent in RT (-). **C.** Sanger sequencing **c**hromatogram showing back-spliced junction of circRNAs amplified using divergent RT-PCR. **D.** Bar plot showing levels of candidate circRNAs and control mRNAs after RNase R degradation, measured by RT-qPCR. **E.** Bar plot depicting differential expression of the five circRNAs during myoblast differentiation, quantified using RT-qPCR. **F.** Schematic showing light monosome and heavy polysome fractions carrying free non-translating and ribosome-bound actively translating circRNAs, respectively (left). Bar graph depicting the distribution of candidate circRNAs in monosome and polysome of C2C12 myotubes by RT-qPCR (right). Data shown in panels D-F represent the mean ±SEM of at least three biological replicates, and * indicates significance with a p-value <0.05.

### *CircSmad1* is a conserved myogenesis-specific circRNA

Among the polysome-associated circRNAs identified in mouse skeletal muscle, we focused on *circSmad1*, which is abundantly expressed across diverse tissues/organs and reported as a potential protein-coding circRNA in several databases, including CIRCpedia v3, circAtlas 3.0, circSC, and TransCirc (**Supplementary Table S4**) [36, 58–60]. *CircSmad1* is back-spliced from the second exon of the Smad1 gene and harbours an open-reading frame encompassing the whole circRNA (**Supplementary Figure S1,** Figure 3A). *CircSmad1* homology was predicted across various mammals including humans and mice. Orthologs of c*ircSmad1* across thirteen different species showed high sequence coverage and identity, indicating evolutionarily conserved biogenesis and function of *circSmad1* (**Supplementary Table S5, Figure S2,** Figure 3B) [36]. Expression of the human *circSMAD1* in differentiating HSMM cells was confirmed by divergent RT-qPCR, followed by Sanger sequencing to validate the back-spliced junction sequence (Figure 3C**,D**). RT-qPCR indicated that *circSmad1* was significantly upregulated in myotubes compared to myoblasts in both mice and human skeletal muscle cells, suggesting myogenesis-specific expression of *circSmad1* (Figure 3E). Further, we tested the subcellular distribution of *circSmad1* in differentiating C2C12 cells and found that *circSmad1* was predominantly localized in the cytoplasm, supporting its availability for ribosomal association and translation into polypeptide (Figure 3F). We also observed that *circSmad1* showed resistance to the RNase R digestion in C2C12 cells, confirming the covalently closed structure of *circSmad1* (Figure 2D). We further determined the stability and half-life of *circSmad1* in C2C12 cells treated with Actinomycin D, a potent transcription inhibitor. We found that after blocking transcription, *circSmad1* exhibited robust cellular levels up to eight hours, compared to linear *cMyc* which showed rapid decay within an hour, implying enhanced longevity of *circSmad1* in skeletal muscle cells (Figure 3G).

**Figure 3:**
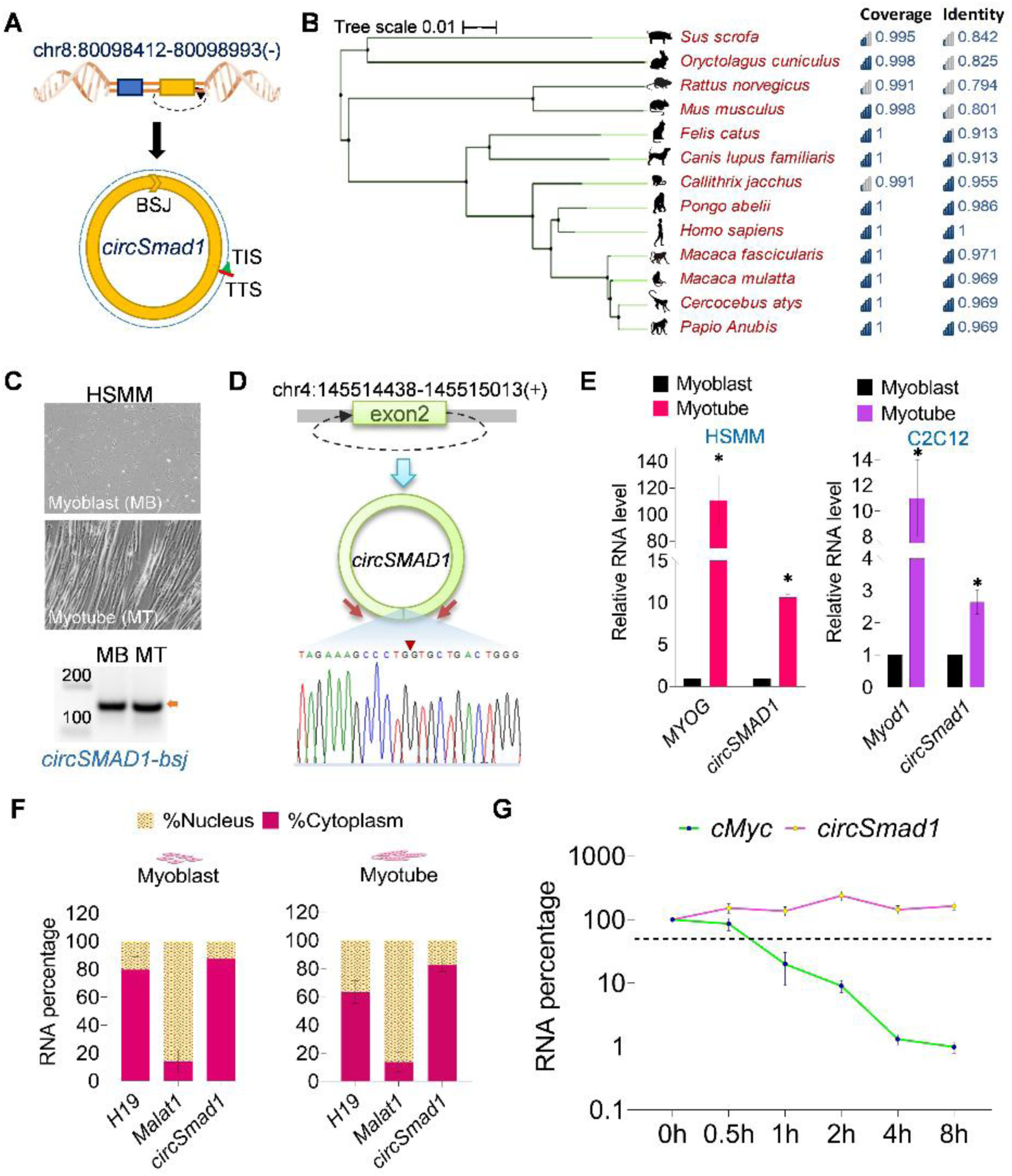
Characterization of *circSmad1* in skeletal muscle cells. **A.** Back-splicing of exon 2 of the *Smad1* gene generates circular *Smad1* (*circSmad1)* containing a junction-crossing ORF. **B.** Phylogenetic tree demonstrating conservation of *circSMAD1* in thirteen mammalian species. **C.** Representative Phase contrast images of HSMM myoblast and myotube (top), agarose gel showing *circSMAD1* back-spliced junction (bsj) band from RT-PCR in HSMM cells (bottom). **D.** Illustrating back-splicing of *circSMAD1* human homolog (top), chromatogram showing *circSMAD1-*bsj confirmed by Sanger sequencing (bottom). **E.** Bar plots showing RT-qPCR expression profile of *circSmad1* during differentiation of myoblasts to myotubes in both HSMM (left) and C2C12 (right) cells, normalised to *18S* rRNA, and *Gapdh* internal control mRNAs, respectively. **F.** Histogram depicting the level of *circSmad1* in the cytoplasm and nucleus of differentiating C2C12 cells by RT-qPCR. *H19* (cytosolic) and *Malat1* (nuclear) were used as marker genes. **G.** Line plot showing half-life of *circSmad1* compared to *cMyc* mRNA in Actinomycin D-treated C2C12 cells, using RT-qPCR. Data represented in panels E-G represent the mean ± SEM of at least three biological replicates, and * indicates significance with a p-value of <0.05.

### Rolling translation of m^6^A-enriched *circSmad1*

*CircSmad1* was predicted to contain an ORF spanning the entire 581 nt circRNA sequence with partially overlapping initiation (A<u>UG</u>) and termination (<u>UG</u>A) codons (**Supplementary Figure S1,** Figure 3A). To determine the protein-coding ability of the putative *circSmad1-*cORF in skeletal muscle cells, we tested the ribosomal association pattern of *circSmad1* during polysome fractionation in C2C12 cells. We detected a minor population of *circSmad1* in the light and heavy polysome fractions, while the majority (> 80%) of the circRNA remained in the monosome or unbound fractions. Although small, a relatively significant polysome association was observed in *circSmad1* in comparison to both *H19* lncRNA (used as negative control) and *circZfp609,* an experimentally verified polysome-bound circRNA (used as positive control) [20]. The association of *circSmad1* with polysomes was confirmed in mouse skeletal muscle cells (Figure 4A**,B**). It has been reported that circRNAs utilise m^6^A motifs to recruit eukaryotic initiation complex (YTHDF3/eIF4G2), and initiate protein translation in a cap-independent fashion [61]. We used the SRAMP webserver to predict the presence of m^6^A sites in *circSmad1* as well as the secondary structure with all possible m^6^A sites (Figure 4C**,D**) [41]. We also cross-referenced *circSmad1* in the m6A2Circ database and collected MeRIP-seq and WER RIP-seq evidence supporting m^6^A-*circSmad1* association in diverse tissues (**Supplementary Table S6**) [40]. We determined the level of N6-methyladenosine modification in *circSmad1* using methylated RNA immunoprecipitation (MeRIP) followed by RT-qPCR in C2C12 cells (Figure 4E). We found significant enrichment of *circSmad1* in RIP with anti-m^6^A antibody compared to the control IgG in C2C12 cells (Figure 4F). Altogether, the MeRIP study indicated that *circSmad1* contains m^6^A modification and may act as a template for cap-independent translation in mouse skeletal muscle.

**Figure 4:**
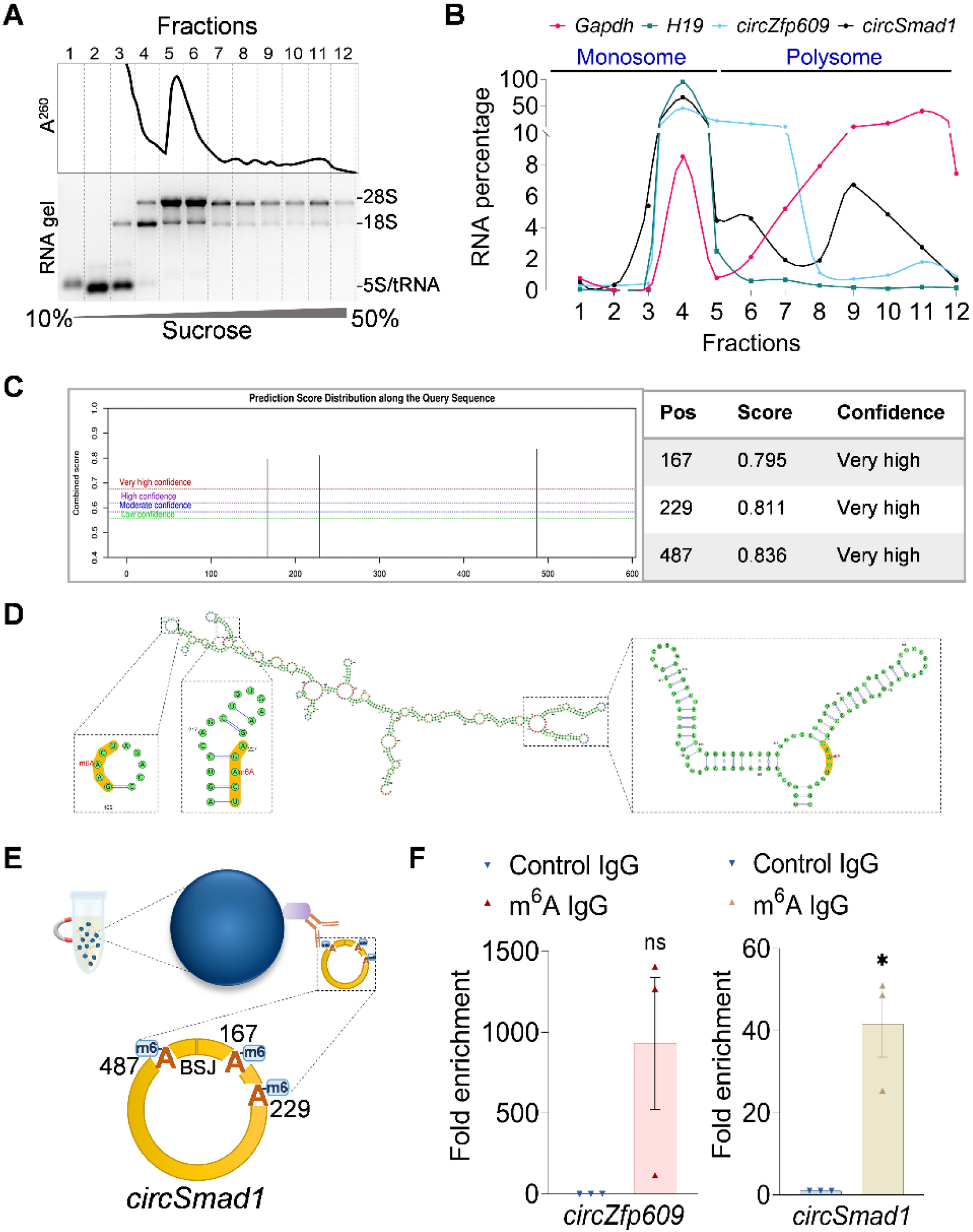
Study of cap-independent translation of m^6^A-enriched *circSmad1* in C2C12 cells. **A.** Representative polysome fractionation graph of C2C12 myotube (top) and detection of total RNA purified from each of the twelve sucrose gradient fractions, resolved on a 1.2% MOPS denaturing gel (bottom). **B.** Line plot showing the distribution of *Gapdh* mRNA, *H19* lncRNA, *circZfp609*, and *circSmad1* circRNAs across monosome (F1-F5) and polysome (F6-F12) fractions in CHX-treated myotubes by RT-qPCR. **C.** Prediction of m^6^A sites in *circSmad1* mature sequence using the SRAMP online tool. **D.** Secondary structure of *circSmad1* predicted by RNAfold, showing m^6^A sites at three different positions. **E.** Illustration of immunoprecipitation of methylated *circSmad1* RNA (MeRIP). **F.** Bar plot showing enrichment levels of *circZfp609* (positive control) (left) and *circSmad1* (right) in the m^6^A pull-down compared to control IgG, measured using RT-qPCR. Data shown in panel F represent the mean ± SEM of three biological replicates, and * indicates significance with a p-value <0.05.

### Downregulation of *circSmad1* impairs differentiation of C2C12 myoblasts

Earlier, we showed that *circSmad1* exhibits greater expression upon differentiation in both C2C12 and HSMM cells by RT-qPCR assay (Figure 3E), indicating that *circSmad1* plays a functional role in myogenesis. We used locked nucleic acid (LNA) antisense oligo Gapmer targeting the *circSmad1* back-spliced junction as the *circSmad1*-Gapmer, while non-targeting LNA Gapmer was used as the control Gapmer. To better understand the function of *circSmad1* in skeletal muscle, we studied the effect of *circSmad1* silencing in C2C12 myoblasts differentiated for four days after transfection with *circSmad1*-Gapmer (Figure 5A). We confirmed that the *circSmad1* level was effectively downregulated in the *circSmad1*-Gapmer samples relative to the control Gapmer (Figure 5B). Interestingly, we observed that myotube formation was visibly impaired and halted at the cell fusion step in the *circSmad1*-Gapmer group, but not in the control group (Figure 5C). Additionally, staining with Jenner-Giemsa clearly indicated reduced myotube density and fusion in the *circSmad1*-Gapmer compared to the control (Figure 5C). Next, we evaluated the effect of *circSmad1* silencing on the levels of myogenesis regulatory genes, five days post-transfection in C2C12 myotubes.

**Figure 5:**
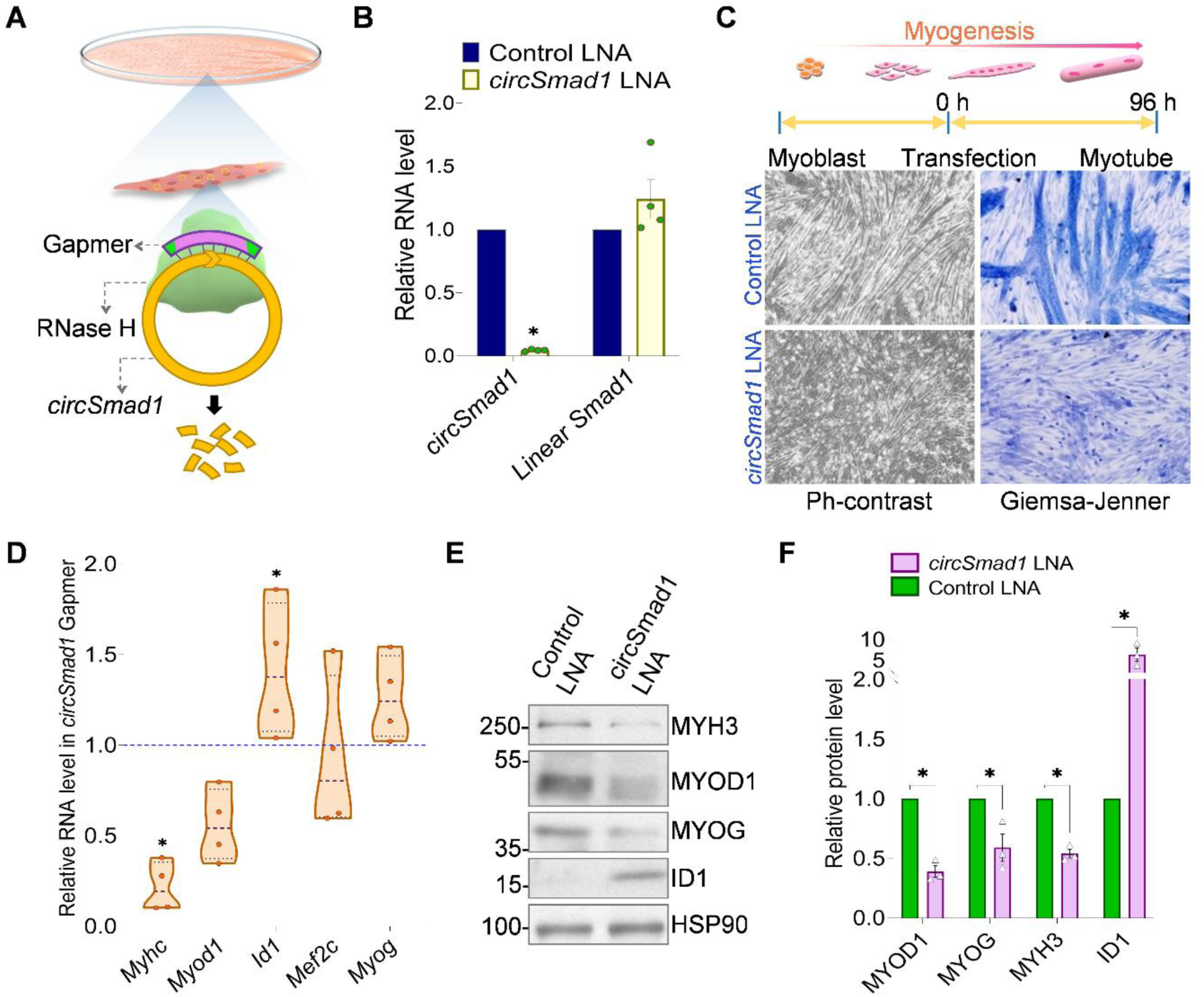
Effect of *circSmad1* silencing on myogenesis in C2C12 cells. **A.** Illustration of *circSmad1*-junction specific LNA Gapmer and RNase H-mediated degradation of *circSmad1* in cultured myotubes. **B.** Histogram showing the level of *circSmad1* and cognate *Smad1* mRNA normalised to *18S* rRNA in *circSmad1-*Gapmer and control Gapmer groups using RT-qPCR. **C.** Representative microscopic images showing reduced myotube formation in *circSmad1-*Gapmer compared to the control group in 4-day differentiated C2C12 myotubes. **D.** Truncated violin plot showing relative expression of myogenic regulatory genes normalised to *18S* rRNA in *circSmad1*-Gapmer myotubes compared to control by RT-qPCR. **E.** Representative western blot images showing bands specific to MRFs in control and *circSmad1*-Gapmer cells. **F.** Densitometry estimation of myogenesis regulatory proteins normalised to HSP90 is represented in the bar plot. Data shown in panels B, D, and F represent the mean ± SEM of at least three biological replicates, and * indicates significance with p-value <0.05.

We found that the expression of Myod1 and Myhc was reduced in *circSmad1*-silenced cells compared to the control, as confirmed by both RT-qPCR and immunoblotting assays (Figure 5D-F). Interestingly, immunofluorescence staining of MRFs in transfected C2C12 myotubes further confirmed the downregulation of MYOG and MYH3 in *circSmad1*-Gapmer group compared to control, while MYOD1 showed a higher cytoplasmic localization upon *circSmad1* silencing (**Supplementary Figure S3**). Overall, our findings showed that knockdown of *circSmad1* downregulates key myogenesis regulatory genes, thereby inhibiting myoblast differentiation and confirmed that *circSmad1* plays an essential role in promoting myogenesis in mice.

### CircSmad1-194aa acts as a nuclear factor mimicking the SMAD1-MH1 domain

Supporting evidence collected from ORF prediction, species conservation, polysome profiling and MeRIP-qPCR assay of *circSmad1* strongly suggests the translation of *circSmad1* into a 194 aa peptide – circSmad1-194aa in C2C12 skeletal muscle cells (**Supplementary Figure S1 and S2,** Figure 2F**, 3B, and 4F)**. Sequence homology comparison of circSmad1-194aa across different mammalian species confirmed that the circSmad1-194aa is highly conserved and may have an evolutionarily retained function (**Supplementary Table S5, Figure S4**). Based on multi-omics evidence curated in riboCIRC v1, the *circSmad1* junction-specific peptide was detected in mouse mass spectrometry dataset using MaxQuant software (**Supplementary Figure S5**) [24, 48, 49]. Furthermore, we investigated the intracellular properties of circSmad1-194aa in C2C12 cells transfected with the pCMV6-circSmad1 vector (Figure 6A). We first confirmed the overexpression of FLAG-tagged circSmad1-194aa in C2C12 cells harvested two days post-transfection. Using anti-FLAG western blotting, we detected a clear band at the expected size (∼24 kDa) in the pCMV6-circSmad1 group, while no band was found in the control pcDNA3 group (Figure 6B). Notably, we found the circSmad1-cORF is composed of an AUG initiation site shared with the host gene, and the cORF ends at a unique stop site due to rolling-circle translation, suggesting a possible sequence overlap between the host SMAD1 and circSmad1-194aa (Figure 3A). Pairwise sequence alignment showed an absolute identity of the circSmad1-194aa to the MAD homology 1 (MH1) domain present near the N-terminus of SMAD1 (**Supplementary Figure S6,** Figure 6C). Similarly, functional site prediction using InterProScan indicated the presence of the conserved MH1 domain at the N-terminal site of circSmad1-194aa (**Supplementary Figure S7**) [45]. Next, we used the RCSB PDB webserver and compared the circSmad1-194aa structure (predicted using AlphaFold2 Colab) with pdb_00003kmp, a publicly available crystal structure of SMAD1-MH1/DNA complex (**Supplementary Figure S8)** [43, 46, 47]. We found that circSmad1-194aa was highly similar to the DNA-binding MH1 domain with a root-mean-square deviation (RMSD) of 1.18, implying a close functional resemblance (Figure 6D).

**Figure 6:**
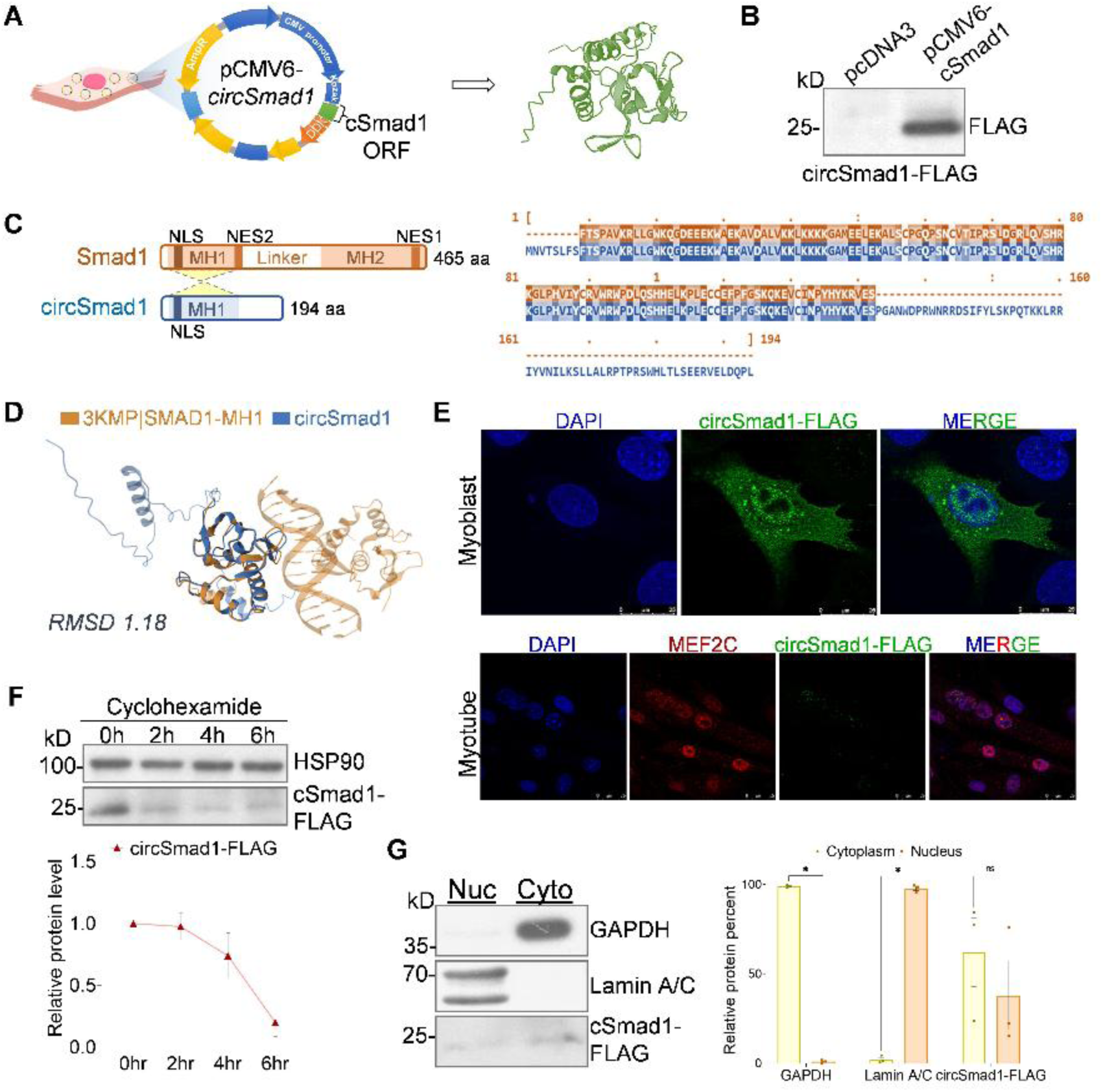
Characterization of circSmad1-194aa in C2C12 cells. **A.** Schematic showing vector map of pCMV6-circSmad1-FLAG expression vector (left) and predicted structure of the circSmad1-194aa protein using AlphaFold v2 (right). **B.** Representative western blotting image showing signal specific to FLAG-tagged circSmad1-194aa present in pCMV6-circSmad1 and absent in the control pcDNA3 sample. **C.** Sequence comparison showing a common MH1 domain structure present in the circSmad1-194aa and host SMAD1, as depicted in the illustration (left) and pair-wise alignment (right) using BLASTp. **D.** Structure homology between circSmad1-194aa and the SMAD1-MH1 domain was demonstrated using the RCSB PDB webserver. **E.** Representative immunofluorescence staining images acquired by confocal microscopy showing subcellular localization of circSmad1-194aa (green) and MEF2C (red) in murine myoblasts (top) and myotubes (bottom) with nuclei counterstained with DAPI (blue). **F.** Representative immunoblotting images showing levels of circSmad1-194aa in C2C12 cells at different time points after cyclohexamide treatment (top) and densitometry quantification depicted as a line plot (bottom). **G.** Representative western blotting images detecting GAPDH, Lamin A/C and circSmad1-194aa across subcellular fractions of C2C12 cells (left) and relative levels shown as bar chart, estimated by densitometry (right). Data shown in panels F and G represent the mean ± SEM of at least three biological replicates, and * indicates significance with a p-value <0.05.

To visualize circSmad1-194aa in the subcellular locations, we performed an immunofluorescence staining with anti-FLAG antibody in differentiating C2C12 cells. We detected circSmad1-194aa signal (green) in both the cytoplasm and the nucleus of C2C12 myoblasts, 48 h post-transfection (Figure 6E**, top**). In contrast, fluorescence signal specific to circSmad1-194aa (green) was only detected in the nuclei of C2C12 myotubes, along with MEF2C (red), a late myogenic factor (Figure 6E**, bottom**). These findings showed that the circSmad1-194aa undergoes active nuclear localization during myoblast differentiation in C2C12 cells, and it may function as a transcription factor in myogenesis. Previous studies have reported that muscle-specific transcription factors such as Myod1 and Myog exhibit a short half-life of nearly one hour [62, 63]. We assessed the stability of circSmad1-194aa in transfected C2C12 cells using cyclohexamide chase assay. We detected a gradually declining signal of circSmad1-194aa till 6 h post-translation inhibition, indicating a robust half-life of circSmad1-194aa, further corroborating its function as a transcription factor in differentiating C2C12 cells (Figure 6F). Consistent with the IF results, we demonstrated the subcellular distribution of circSmad1-194aa in C2C12 cells, 48h post-transfection, by immunoblotting with anti-FLAG antibody, indicating localization of circSmad1-194aa in both the cytoplasmic and nuclear fractions of C2C12 myoblasts (Figure 6G). Interestingly, we detected a significantly higher signal of circSmad1-194aa in the insoluble fraction compared to the soluble fraction of the nucleus in C2C12 myoblasts (**Supplementary Figure S9**). Altogether, the findings suggested that circSmad1-194aa is not freely nucleoplasmic but chromatin-associated and undergoes myogenesis-induced nuclear translocation due to the presence of a nuclear localization signal (NLS), as part of the MH1 domain (Figure 6C).

### Competitive binding of circSmad1-194aa to the *Id1-*BRE promotes myogenesis

Several studies established that Bone Morphogenetic Protein (BMP) signalling inhibits myogenesis in mice [64]. BMP-2 stimulation recruits SMAD1-SMAD4 heterocomplexes in C2C12 myoblasts, translocating to the nucleus to bind a BMP-responsive element (BRE) ∼1 kb upstream of the *Id1* transcription start site, eliciting *Id1* expression (Figure 7A). Id1 sequesters E-proteins, which inhibits basic helix-loop-helix (bHLH) MRFs such as Myod1 and Myogenin, effectively blocking myogenesis [54]. Structural homology and IF results were consistent with the hypothesis that circSmad1-194aa resembles the DNA-binding MH1 domain, which is highly conserved in MAD family proteins, and binds to the BRE near the proximal promoter region of the *Id1* gene in myocytes [65, 66]. We asked whether circSmad1-194aa, which contains an intact MH1 domain, may occupy the *Id1*-BRE promoter site, rendering it unavailable for any physical interaction with the SMAD1-SMAD4 complex. We further demonstrated the interaction between circSmad1-194aa and *Id1*-BRE in C2C12 cells transfected with the circSmad1-194aa overexpression vector using the ChIP-qPCR assay (Figure 7B). We found that the *Id1-*BRE showed greater enrichment levels relative to input in the anti-FLAG group, while a negligible amount was detected in the control IgG. We used *18S* rRNA sequence as a negative control, which showed no enrichment relative to the input in both the anti-FLAG and anti-IgG, confirming specific binding of circSmad1-194aa to the *Id1*-BRE locus in C2C12 chromatin (**Supplementary Table S3,** Figure 7C). We further investigated the association between circSmad1-194aa and *Id1*-BRE in vitro using a biotinylated oligo immunoprecipitation assay with a 40bp 3’biotin-labelled *Id1*-BRE dsDNA probe in C2C12 lysate, followed by western blotting with anti-FLAG antibody (**Supplementary Table S3**, Figure 7D**,E**). We detected a significant increase in the amount of circSmad1-194aa in *Id1*-BRE compared to the control probe, suggesting that the circSmad1-194aa binds to the *Id1-*BRE with greater affinity (Figure 7F). Altogether, the ChIP and biotinylated dsDNA pull-down results confirmed that circSmad1-194aa interacts with the BRE region of *Id1* in C2C12 cells. Interestingly, we also demonstrated that silencing of *circSmad1* led to a significant increase in the level of *Id1* in C2C12 cells (**Supplementary Figure S3,** Figure 5D-F), suggesting that the protein-coding *circSmad1* regulates myogenesis via circSmad1-194aa, which acts as a transcription modulator of *Id1* in C2C12 muscle cells (Figure 8).

**Figure 7.**
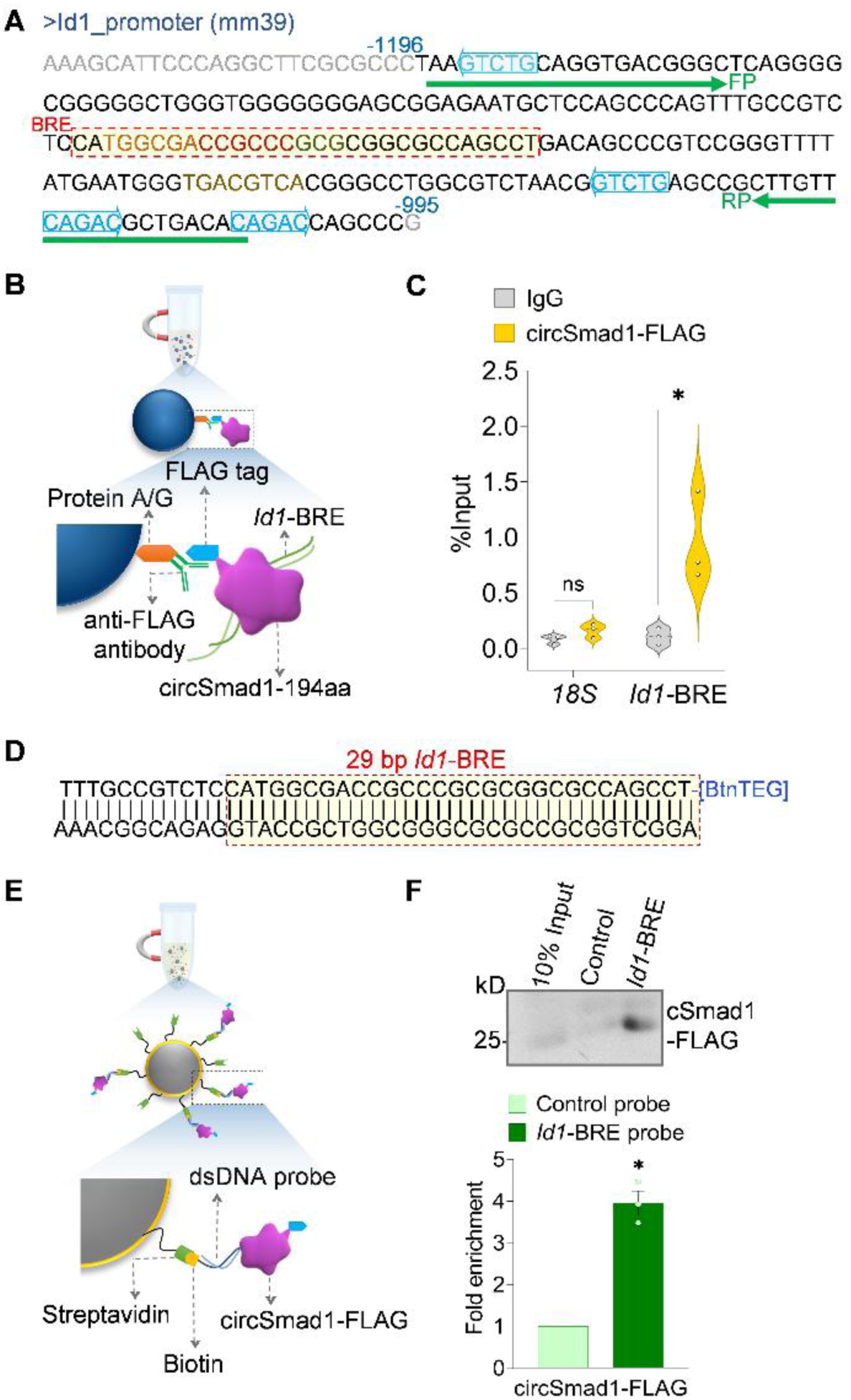
Interaction of circSmad1-194aa with *Id1* promoter in C2C12 cells. **A.** Consensus sequence of a 225bp region of the mouse *Id1* promoter, including CAGAC boxes (blue) and BMP responsive element (BRE) highlighted in a dotted red box. The green arrows correspond to primers targeting *Id1*-BRE. **B.** Schematic of chromatin immunoprecipitation (ChIP) of FLAG-tagged circSmad1-194aa in C2C12 lysate. **C**. Violin plot showing the level of *Id1-*BRE in circSmad1-FLAG and control IgG estimated by ChIP-qPCR assay, with *18S* rRNA used as a negative control. **D.** Annealed double-stranded sequence of 3’ biotinylated *Id1*-BRE probe (BRE region marked in box). **E.** Schematic of streptavidin**-**biotin oligo immunoprecipitation of FLAG-tagged circSmad1-194aa in C2C12 lysate. **F.** Representative immunoblotting image showing signal of circSmad1-FLAG in *Id1*-BRE compared to the control and input (top), and its densitometry estimation plotted as a bar graph (bottom). Data shown in panels C and F represent the mean ± SEM of at least three biological replicates, and * indicates significance with a p-value <0.05.

**Figure 8:**
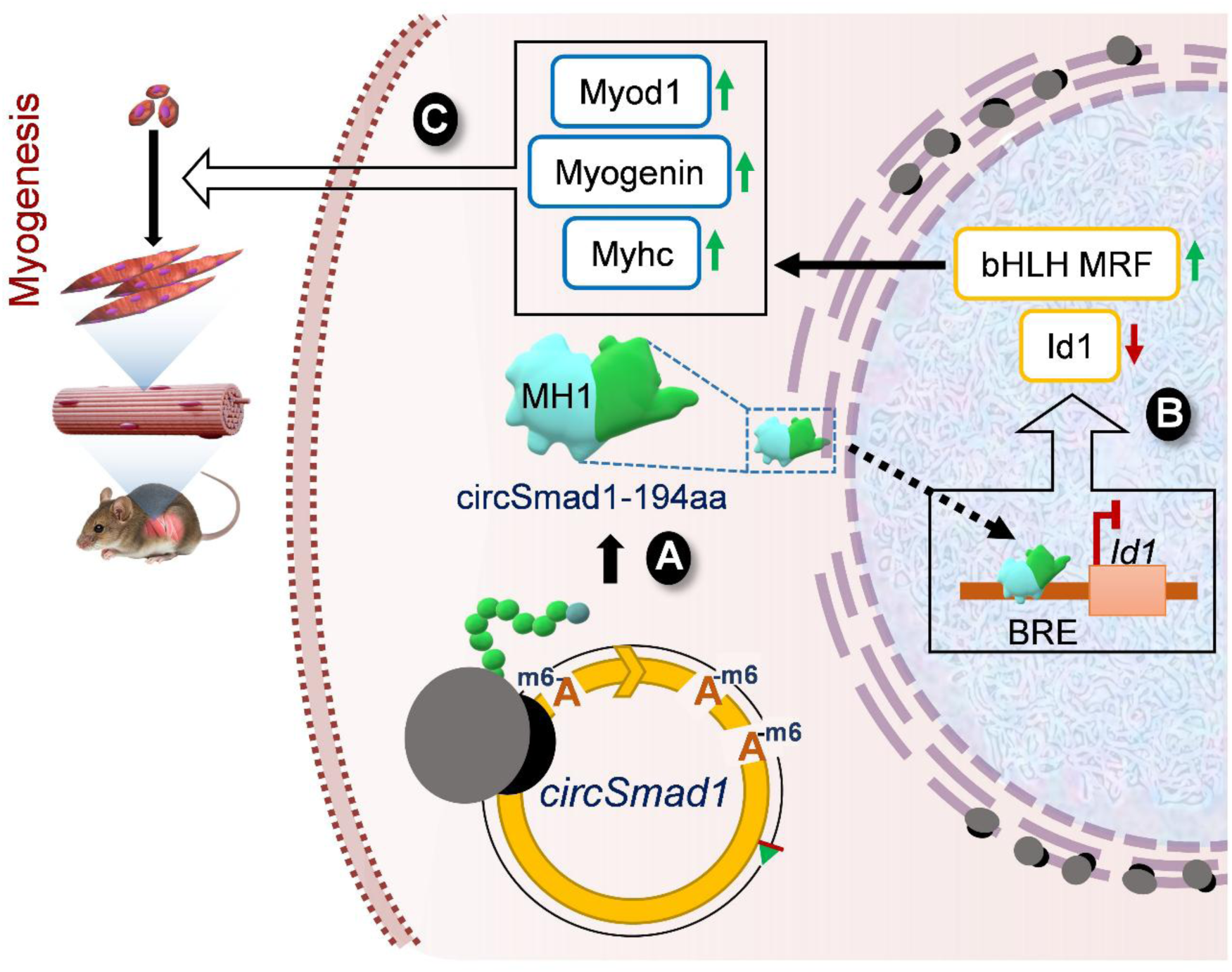
Role of circSmad1-194aa in murine myogenesis. **A.** *CircSmad1* contains a junction-spanning ORF and multiple m6A sites, which recruit polysomes and undergo cap-independent translation into a 194aa polypeptide called circSmad1-194aa. **B.** CircSmad1-194aa contains a conserved DNA-binding MH1 domain, which mediates its nuclear translocation and competitive binding at the *Id1*-BRE promoter site, leading to downregulation of *Id1*. **C.** Reduced expression of *Id1* derepresses the activity of basic HLH myogenesis regulatory factors (Myod1 and Myogenin), inducing myogenesis in mouse C2C12 myoblasts.

## DISCUSSION

CircRNAs are an emerging class of RNA molecules that indirectly regulate gene expression across diverse pathophysiological conditions by interacting with miRNAs, RNA-binding proteins, RNA polymerase II, and mRNAs [9–11]. Interestingly, recent studies have shown that circRNAs can be translated into peptides through cap-independent mechanisms, highlighting a direct function of protein-coding circRNAs [17, 18, 27]. Typically, circRNAs are back-spliced into a closed-loop structure that can undergo rolling-circle translation via cap-independent initiation at internal ribosome entry sites (IRESs), N6-methyladenosine (m^6^A), and exon junction complex (EJC) [19, 61]. In 2017, Legnini *et al* first demonstrated that *circZNF609* harbours an IRES that recruits polysomes and enables its translation into a protein with a functional role in myogenesis [20]. More recent studies have identified different translatable circRNAs in skeletal muscle across species, including *circFAM188B* in chicken, *circDdb1* in mice, and *circNEB* in cattle [21–23]. Skeletal muscle provides a favourable environment for circRNA accumulation and translation, as these highly stable RNA molecules function in well-differentiated tissues and organs [20, 67]. Existing mass spectrometry pipelines remain biased against short, low-abundance peptides, limiting the detection of microproteins relevant to myogenesis. Overcoming such challenges will require integrated strategies that combine deep ribosome profiling, circRNA discovery, proteogenomics, mass spectrometry, and functional validation in physiologically relevant muscle models.

In this study, we combined polysome RNA-seq in C2C12 cells with riboCIRC v1, which compiles Ribo-seq, MS/MS, m^6^A and IRES-based evidence for circRNA translation, and identified hundreds of putative protein-coding circRNAs in mouse skeletal muscle (Figure 1) [24]. Polysome association is a robust indicator of circRNA translation, particularly in terminally differentiated tissues such as skeletal muscle [14, 18, 68]. We validated the upregulation of two polysome-associated circRNAs – *circSmad1* and *circH6pd* with high coding potential in C2C12 myotubes using divergent RT-qPCR, back-spliced junction sequencing, and RNase R resistance assay (Figure 2). Among the polysome-associated circRNAs, we discovered circular *Smad1* (*circSmad1*), which originates from back-splicing of exon 2 of the Smad1 gene and is publicly reported to be abundantly expressed in various tissues and disease conditions [36, 58–60]. Consistent with previous studies, multiple-sequence alignment of *circSmad1* with cross-species homologs curated in CIRCpedia (v3) revealed strong evolutionary conservation, a common feature of functionally relevant circRNAs (Figure 3) [36]. Divergent RT-qPCR analysis confirmed that *circSmad1* is enriched in mouse/human myotubes and is predominantly localized to the cytoplasm, similar to the characteristics of protein-coding circRNAs involved in myogenesis (Figure 3) [20].

Furthermore, actinomycin D chase assays in C2C12 cells demonstrated that *circSmad1* is highly stable with a half-life exceeding 8 h, supporting its potential for efficient cap-independent translation due to the inherent stability of circRNAs relative to linear counterparts (Figure 3). ORF prediction identified a junction-spanning open reading frame covering the full *circSmad1* sequence, initiating at the canonical host AUG and encoding a 194 aa peptide (circSmad1-194aa) (Figure 3). Cross-species ORF prediction indicated a high degree of conservation of circSmad1-194aa, consistent with examples of actively translated circRNAs [20, 23]. Apart from ORF, circRNAs depend on certain structural features, such as IRES and m^6^A, to recruit the translation pre-initiation machinery in the absence of the 5’ cap. Reportedly, m^6^A modification is highly prevalent in all forms of RNA, and it effectively drives circRNA cap-independent translation through the YTHDF3/eIF4G2/eIF3A complex [19, 40, 61]. MeRIP-qPCR analysis showed significant enrichment of m⁶A on *circSmad1* in C2C12 cells, comparable to the protein-coding *circZfp609*, suggesting that m⁶A modification may facilitate *circSmad1* translation in mouse myocytes (Figure 4). Importantly, *circSmad1* junction-specific peptide was detected in mouse proteome raw data previously reported in riboCIRC (v1) as MS evidence for *circSmad1*, providing direct support for bona fide *circSmad1* translation in mice [24].

Skeletal muscle differentiation depends on the fusion of myoblasts into multinucleated myotubes, which serves as a critical step in the process of myogenesis [3, 4, 57]. In this study, we demonstrated that *circSmad1* encodes a peptide (circSmad1-194aa), which promotes mouse myoblast differentiation in vitro. Notably, knockdown of *circSmad1* in C2C12 cells led to aberrant myoblast fusion and reduced myotube formation, accompanied by decreased Myod1, Myog, and Myhc levels, underscoring the functional importance of *circSmad1* in myogenesis. Interestingly, silencing of *circSmad1,* which encodes circSmad1-194aa, significantly elevated *Id1* levels, suggesting a regulatory role of circSmad1-194aa in *Id1* expression (Figure 5). Structure homology showed that circSmad1-194aa closely resembles the Smad1/MH1 domain, which specifically binds the BMP-responsive element (BRE) of *Id1* (Inhibitor of DNA binding I) in C2C12 cells [54]. Immunofluorescence revealed that circSmad1-194aa is primarily localized to the nucleus in myotubes, consistent with the DNA-binding function of circSmad1-194aa in controlling *Id1* expression (Figure 6). Additionally, the cycloheximide chase assay revealed that circSmad1-194aa is a relatively stable peptide with a half-life exceeding two hours, a property comparable to the well-known basic helix-loop-helix transcription factors in myogenesis, such as Myod1 and Myog (Figure 6) [62, 63].

*Id1* is a well-established inhibitor of myogenesis that sequesters E-proteins and suppresses the activity of bHLH MRFs, including Myod1 and Myogenin [69, 70]. Its expression is temporally regulated, with high levels in myoblasts that decline during differentiation, a pattern essential for myotube formation [71]. The *Id1* promoter contains a 29bp GC-rich element (BRE) that recruits Smad1-Smad4 in response to BMP-2, resulting in increased *Id1* levels that inhibit myogenic differentiation [54]. SMAD complexes recognise GC-rich and CAGAC motifs via the MH1 domain [46]. Our results suggest that the interaction between circSmad1-194aa and *Id1*-BRE counteracts BMP-mediated *Id1* induction, relieving repression of *Myod1*, and promoting myogenesis, as confirmed by ChIP and biotin-oligo pull-down analysis (Figure 7).

Our findings align with previous studies suggesting that BMP ligands inhibit C2C12 differentiation and promote osteogenic fate, whereas attenuation of BMP signalling enhances myogenesis [72]. BMP signalling also influences adult muscle mass and hypertrophy, in contrast to TGF-β family members such as myostatin, which suppresses muscle growth [73]. Cap-independent translation of *circSmad1* produces circSmad1-194aa, a truncated Smad1 variant lacking the MH2 transactivation domain. Functionally, circSmad1-194aa resembles inhibitory Smads (Smad6/7), which hinder BMP signalling by preventing Smad1-Smad4 complex formation, but it operates through a distinct mechanism involving direct competition for DNA binding sites [74]. This is consistent with structural studies showing that the MH1 domain can bind BRE motifs independently of MH2, which is finely regulated by conformation, cofactors, and DNA motifs [75]. The biological relevance of the circSmad1-194aa mechanism is further supported by a study showing that isolated Smad1 MH1 retains DNA-binding capacity but lacks full transcriptional activity in mesenchymal progenitors, behaving as a context-dependent modulator rather than an activator [72]. Our findings acknowledge a previously unrecognised node of SMAD1 regulation and highlight circRNA-derived proteins as potent regulators of muscle fate with potential therapeutic relevance.

In summary, we demonstrated that *circSmad1* encodes circSmad1-194aa, a naturally occurring polypeptide containing MH1 domain that acts as a negative factor restraining BMP-mediated *Id1* activation and thereby promoting *Myod1*-driven myogenic differentiation (Figure 8). However, this study has a few limitations that can be addressed in follow-up studies. The protein-coding circRNAs in C2C12 were identified from polysome-associated RNA-seq data only. Ribo-seq in differentiating mouse C2C12 and human HSMM cells need to be performed to find high-confidence ribosome-associated circRNAs in both species. The RNA-seq analysis was performed only with the CIRCexplorer2 pipeline. Other pipelines, along with analysis of the full-length sequence of circRNAs, may identify novel circRNAs and their splice variants associated with translating polysomes in muscle cells [27]. Most of the conclusions are based on experiments performed in cultured cells and should be extended to animal models for a better understanding of the physiological relevance of the *circSmad1*-194aa peptide in skeletal muscle. Although we established a pro-myogenic role for circSmad1-194aa in vitro, its application in muscle regeneration and therapy remains to be explored.

These findings expand our understanding of circRNA function by highlighting the role of protein-coding circRNAs in skeletal muscle, a largely unexplored area. In conclusion, we identified a *circSmad1*-encoded peptide that regulates *Id1* expression to promote myogenesis, and further emphasise circRNA-derived peptides as promising tools for modulating muscle development, ageing, and regeneration.

## ACKNOWLEDGEMENTS

The authors thank Regional Centre for Biotechnology (RCB), Faridabad for PhD registration of Tanvi Sinha and Swarnava Dutta. We thank our lab colleagues Dishanee Santra, Gaurhari Sahoo and Preeti Pranjya Mohapatra for the helpful discussion and for proofreading the manuscript. In addition, we thank the ILS Central Instrumentation Facility (CIF) for helping us with the NGS and Confocal Imaging experiments, funded by Department of Biotechnology (BT/INF/22/SP28293/2018).

## AUTHOR CONTRIBUTIONS

Tanvi Sinha: Conceptualization, Formal analysis, Methodology, Visualization, Validation, Writing—original draft, Writing—review & editing. Swarnava Dutta: Formal analysis, Methodology, Visualization, Validation, Writing—review & editing. Punit Prasad: Resources, Formal analysis, Methodology, Supervision, Writing—review & editing. Amaresh C. Panda: Conceptualization, Fund acquisition, Resources, Supervision, Formal analysis, Writing—original draft, Writing—review & editing..

## SUPPLEMENTARY DATA

Supplementary Data are available Online.

## DATA AVAILABILITY

All the data generated in this study are included in the main text or Supplementary data. The RNA-seq data generated in this study were deposited in the Indian Nucleotide Data Archive (INDA) with International Nucleotide Sequence Database Collaboration (INSDC) Accession number PRJEB108758.

## REFERENCES

1. Frontera, W.R. and J. Ochala, Skeletal muscle: a brief review of structure and function. Calcif Tissue Int, 2015. 96(3): p. 183–95.

2. Wolfe, R.R., The underappreciated role of muscle in health and disease. Am J Clin Nutr, 2006. 84(3): p. 475–82.

3. Hernandez-Hernandez, J.M., et al., The myogenic regulatory factors, determinants of muscle development, cell identity and regeneration. Semin Cell Dev Biol, 2017. 72: p. 10–18.

4. Zammit, P.S., Function of the myogenic regulatory factors Myf5, MyoD, Myogenin and MRF4 in skeletal muscle, satellite cells and regenerative myogenesis. Semin Cell Dev Biol, 2017. 72: p. 19–32.

5. Das, A., et al., Circular RNAs in myogenesis. Biochim Biophys Acta Gene Regul Mech, 2020. 1863(4): p. 194372.

6. Hitachi, K., M. Honda, and K. Tsuchida, The Functional Role of Long Non-Coding RNA in Myogenesis and Skeletal Muscle Atrophy. Cells, 2022. 11(15).

7. Wang, J., et al., Effects of microRNAs on skeletal muscle development. Gene, 2018. 668: p. 107–113.

8. Patop, I.L., S. Wust, and S. Kadener, Past, present, and future of circRNAs. EMBO J, 2019. 38(16): p. e100836.

9. Singh, S., T. Sinha, and A.C. Panda, Regulation of microRNA by circular RNA. Wiley Interdiscip Rev RNA, 2023: p. e1820.

10. Das, A., et al., Emerging Role of Circular RNA-Protein Interactions. Noncoding RNA, 2021. 7(3).

11. Singh, S., et al., Global identification of mRNA-interacting circular RNAs by CLiPPR-Seq. Nucleic Acids Res, 2024. 52(6): p. e29.

12. Wright, B.W., et al., The dark proteome: translation from noncanonical open reading frames. Trends Cell Biol, 2022. 32(3): p. 243–258.

13. Posner, Z., I. Yannuzzi, and J.R. Prensner, Shining a light on the dark proteome: Non-canonical open reading frames and their encoded miniproteins as a new frontier in cancer biology. Protein Sci, 2023. 32(8): p. e4708.

14. Patraquim, P., et al., Developmental regulation of canonical and small ORF translation from mRNAs. Genome Biol, 2020. 21(1): p. 128.

15. Dong, X., et al., Small Open Reading Frame-Encoded Micro-Peptides: An Emerging Protein World. Int J Mol Sci, 2023. 24(13).

16. Bonilauri, B. and B. Dallagiovanna, Microproteins in skeletal muscle: hidden keys in muscle physiology. J Cachexia Sarcopenia Muscle, 2022. 13(1): p. 100–113.

17. Sinha, T., et al., Circular RNA translation, a path to hidden proteome. Wiley Interdiscip Rev RNA, 2022. 13(1): p. e1685.

18. Pamudurti, N.R., et al., Translation of CircRNAs. Mol Cell, 2017. 66(1): p. 9–21 e7.

19. Hwang, H.J. and Y.K. Kim, Molecular mechanisms of circular RNA translation. Exp Mol Med, 2024. 56(6): p. 1272–1280.

20. Legnini, I., et al., Circ-ZNF609 Is a Circular RNA that Can Be Translated and Functions in Myogenesis. Mol Cell, 2017. 66(1): p. 22–37 e9.

21. Yin, H., et al., Circular RNA CircFAM188B Encodes a Protein That Regulates Proliferation and Differentiation of Chicken Skeletal Muscle Satellite Cells. Front Cell Dev Biol, 2020. 8: p. 522588.

22. Zhu, X., et al., EIF4A3-Induced Circular RNA CircDdb1 Promotes Muscle Atrophy through Encoding a Novel Protein CircDdb1-867aa. Adv Sci (Weinh), 2024. 11(45): p. e2406986.

23. Huang, K., et al., A Circular RNA Generated from Nebulin (NEB) Gene Splicing Promotes Skeletal Muscle Myogenesis in Cattle as Detected by a Multi-Omics Approach. Adv Sci (Weinh), 2024. 11(3): p. e2300702.

24. Li, H., et al., riboCIRC: a comprehensive database of translatable circRNAs. Genome Biol, 2021. 22(1): p. 79.

25. Velica, P. and C.M. Bunce, A quick, simple and unbiased method to quantify C2C12 myogenic differentiation. Muscle Nerve, 2011. 44(3): p. 366–70.

26. Panda, A.C., J.L. Martindale, and M. Gorospe, Polysome Fractionation to Analyze mRNA Distribution Profiles. Bio Protoc, 2017. 7(3).

27. Das, A., et al., Identification of potential proteins translated from circular RNA splice variants. Eur J Cell Biol, 2023. 102(1): p. 151286.

28. Das, A., et al., A quick and cost-effective method for DNA-free total RNA isolation using magnetic silica beads. Wellcome Open Research, 2023. 8.

29. Chen, S., et al., *fastp: an ultra-fast all-in-one FASTQ preprocessor*. Bioinformatics, 2018. 34(17): p. i884–i890.

30. Dobin, A., et al., STAR: ultrafast universal RNA-seq aligner. Bioinformatics, 2013. 29(1): p. 15–21.

31. Zhang, X.O., et al., Diverse alternative back-splicing and alternative splicing landscape of circular RNAs. Genome Res, 2016. 26(9): p. 1277–87.

32. Quinlan, A.R. and I.M. Hall, BEDTools: a flexible suite of utilities for comparing genomic features. Bioinformatics, 2010. 26(6): p. 841–2.

33. Livak, K.J. and T.D. Schmittgen, Analysis of relative gene expression data using real-time quantitative PCR and the 2(-Delta Delta C(T)) Method. Methods, 2001. 25(4): p. 402–8.

34. Panda, A.C. and M. Gorospe, Detection and Analysis of Circular RNAs by RT-PCR. Bio Protoc, 2018. 8(6).

35. Ratnadiwakara, M. and M.L. Anko, mRNA Stability Assay Using transcription inhibition by Actinomycin D in Mouse Pluripotent Stem Cells. Bio Protoc, 2018. 8(21): p. e3072.

36. Zhai, S.N., et al., CIRCpedia v3: an interactive database for circular RNA characterization and functional exploration. Nucleic Acids Res, 2026. 54(D1): p. D78–D88.

37. Letunic, I. and P. Bork, Interactive Tree Of Life (iTOL): an online tool for phylogenetic tree display and annotation. Bioinformatics, 2007. 23(1): p. 127–8.

38. McWilliam, H., et al., Analysis Tool Web Services from the EMBL-EBI. Nucleic Acids Res, 2013. 41(Web Server issue): p. W597–600.

39. Gruber, A.R., et al., The Vienna RNA websuite. Nucleic Acids Res, 2008. 36(Web Server issue): p. W70–4.

40. Li, Y., et al., m6A2Circ: A comprehensive database for decoding the regulatory relationship between m6A modification and circular RNA. Comput Struct Biotechnol J, 2025. 27: p. 813–820.

41. Zhou, Y., et al., SRAMP: prediction of mammalian N6-methyladenosine (m6A) sites based on sequence-derived features. Nucleic Acids Res, 2016. 44(10): p. e91.

42. Wheeler, D.L., et al., Database resources of the National Center for Biotechnology Information. Nucleic Acids Res, 2008. 36(Database issue): p. D13–21.

43. Jumper, J., et al., Highly accurate protein structure prediction with AlphaFold. Nature, 2021. 596(7873): p. 583–589.

44. Camacho, C., et al., BLAST+: architecture and applications. BMC Bioinformatics, 2009. 10: p. 421.

45. Quevillon, E., et al., InterProScan: protein domains identifier. Nucleic Acids Res, 2005. 33(Web Server issue): p. W116–20.

46. BabuRajendran, N., et al., Structure of Smad1 MH1/DNA complex reveals distinctive rearrangements of BMP and TGF-beta effectors. Nucleic Acids Res, 2010. 38(10): p. 3477–88.

47. Burley, S.K., et al., Updated resources for exploring experimentally-determined PDB structures and Computed Structure Models at the RCSB Protein Data Bank. Nucleic Acids Res, 2025. 53(D1): p. D564–D574.

48. Cox, J. and M. Mann, MaxQuant enables high peptide identification rates, individualized p.p.b.-range mass accuracies and proteome-wide protein quantification. Nat Biotechnol, 2008. 26(12): p. 1367–72.

49. Jones, P., et al., PRIDE: a public repository of protein and peptide identifications for the proteomics community. Nucleic Acids Res, 2006. 34(Database issue): p. D659–63.

50. Yao, Y., et al., METTL3-dependent m(6)A modification programs T follicular helper cell differentiation. Nat Commun, 2021. 12(1): p. 1333.

51. Asakura, A. and N. Kikyo, Immunofluorescence analysis of myogenic differentiation. Methods Cell Biol, 2022. 170: p. 117–125.

52. Wang, J., et al., Cigarette smoking inhibits myoblast regeneration by promoting proteasomal degradation of NPAT protein and hindering cell cycle progression. Curr Res Toxicol, 2024. 6: p. 100161.

53. Holliday, H., A. Khoury, and A. Swarbrick, Chromatin immunoprecipitation of transcription factors and histone modifications in Comma-Dbeta mammary epithelial cells. STAR Protoc, 2021. 2(2): p. 100514.

54. Katagiri, T., et al., Identification of a BMP-responsive element in Id1, the gene for inhibition of myogenesis. Genes Cells, 2002. 7(9): p. 949–60.

55. Das, D., A. Das, and A.C. Panda, Antisense Oligo Pulldown of Circular RNA for Downstream Analysis. Bio Protoc, 2021. 11(14): p. e4088.

56. Tsuji, Y., Optimization of Biotinylated RNA or DNA Pull-Down Assays for Detection of Binding Proteins: Examples of IRP1, IRP2, HuR, AUF1, and Nrf2. Int J Mol Sci, 2023. 24(4).

57. Bentzinger, C.F., Y.X. Wang, and M.A. Rudnicki, Building muscle: molecular regulation of myogenesis. Cold Spring Harb Perspect Biol, 2012. 4(2).

58. Huang, W., et al., TransCirc: an interactive database for translatable circular RNAs based on multi-omics evidence. Nucleic Acids Res, 2021. 49(D1): p. D236–D242.

59. Wu, W., et al., Exploring the cellular landscape of circular RNAs using full-length single-cell RNA sequencing. Nat Commun, 2022. 13(1): p. 3242.

60. Wu, W., F. Zhao, and J. Zhang, circAtlas 3.0: a gateway to 3 million curated vertebrate circular RNAs based on a standardized nomenclature scheme. Nucleic Acids Res, 2024. 52(D1): p. D52–D60.

61. Yang, Y., et al., Extensive translation of circular RNAs driven by N(6)-methyladenosine. Cell Res, 2017. 27(5): p. 626–641.

62. Sun, L., et al., Ubiquitin-proteasome-mediated degradation, intracellular localization, and protein synthesis of MyoD and Id1 during muscle differentiation. J Biol Chem, 2005. 280(28): p. 26448–56.

63. Huang, S.C., et al., Protein 4.1R Influences Myogenin Protein Stability and Skeletal Muscle Differentiation. J Biol Chem, 2016. 291(49): p. 25591–25607.

64. Ono, Y., et al., BMP signalling permits population expansion by preventing premature myogenic differentiation in muscle satellite cells. Cell Death Differ, 2011. 18(2): p. 222–34.

65. Chai, N., et al., Structural basis for the Smad5 MH1 domain to recognize different DNA sequences. Nucleic Acids Res, 2015. 43(18): p. 9051–64.

66. Lopez-Rovira, T., et al., Direct binding of Smad1 and Smad4 to two distinct motifs mediates bone morphogenetic protein-specific transcriptional activation of Id1 gene. J Biol Chem, 2002. 277(5): p. 3176–85.

67. Greco, S., et al., Circular RNAs in Muscle Function and Disease. Int J Mol Sci, 2018. 19(11).

68. van Heesch, S., et al., The Translational Landscape of the Human Heart. Cell, 2019. 178(1): p. 242–260 e29.

69. Lingbeck, J.M., et al., In vivo interactions of MyoD, Id1, and E2A proteins determined by acceptor photobleaching fluorescence resonance energy transfer. FASEB J, 2008. 22(6): p. 1694–701.

70. Trausch-Azar, J.S., et al., Ubiquitin-Proteasome-mediated degradation of Id1 is modulated by MyoD. J Biol Chem, 2004. 279(31): p. 32614–9.

71. AlSudais, H., N. Lala-Tabbert, and N. Wiper-Bergeron, CCAAT/Enhancer Binding Protein beta inhibits myogenic differentiation via ID3. Sci Rep, 2018. 8(1): p. 16613.

72. Ju, W., et al., The bone morphogenetic protein 2 signaling mediator Smad1 participates predominantly in osteogenic and not in chondrogenic differentiation in mesenchymal progenitors C3H10T1/2. J Bone Miner Res, 2000. 15(10): p. 1889–99.

73. Sartori, R., et al., BMP signaling controls muscle mass. Nat Genet, 2013. 45(11): p. 1309–18.

74. Itoh, S. and P. ten Dijke, Negative regulation of TGF-beta receptor/Smad signal transduction. Curr Opin Cell Biol, 2007. 19(2): p. 176–84.

75. Jatzlau, J., et al., Rare but specific: 5-bp composite motifs define SMAD binding in BMP signaling. BMC Biol, 2025. 23(1): p. 79.

