## Supplementary Figure for "Circular Smad1-Encoded Polypeptide Regulates Myogenesis"

### SUPPLEMENTARY TABLES AND FIGURES

**Supplementary Table S1:** Polysome-associated circRNAs in C2C12 cells

**Supplementary Table S2:** Polysome-associated C2C12 circRNAs in riboCIRC v1

**Supplementary Table S3:** Oligonucleotides used in this study

**Supplementary Table S4:** circSmad1 expression profile in other databases

**Supplementary Table S5:** circSmad1 conservation data

**Supplementary Table S6:** circSmad1 data in m6A2Circ database

**Sinha et al, Supplementary Figure S1**

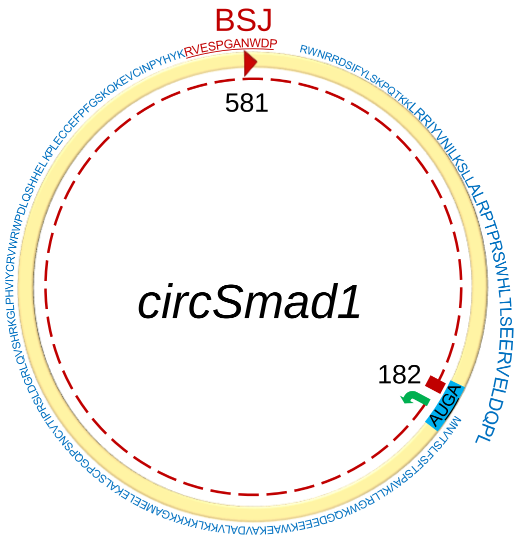


M N V T S L F S F T S P A V K R L L G W
ATGAATGTGACCAGCTTGTTTTCATTCACAAGTCCAGCTGTGAAGAGACTCCTTGGGTGG
K Q G D E E E K W A E K A V D A L V K K
AAACAGGGCGATGAAGAAGAGAAATGGGCAGAGAAAGCTGTGGACGCTTTGGTGAAGAAA
L K K K K G A M E E L E K A L S C P G Q
CTGAAGAAGAAGAAAGGGGCCATGGAAGAGCTGGAGAAGGCCCTGAGCTGCCCTGGACAG
P S N C V T I P R S L D G R L Q V S H R
CCGAGTAACTGCGTCACCATTCCTCGCTCCCTGGATGGCAGGTTGCAGGTGTCCCACCGG
K G L P H V I Y C R V W R W P D L Q S H
AAGGGACTACCTCATGTCATTTATTGCCGTGTGTGGCGCTGGCCCGACCTCCAGAGCCAC
H E L K P L E C C E F P F G S K Q K E V
CATGAACTGAAGCCTCTGGAATGCTGTGAGTTCCCATTTGGTTCCAAGCAGAAGGAGGTC
C I N P Y H Y K R V E S P G A N W D P R
TGCATCAACCCCTACCACTATAAGCGAGTGGAGAGCCCGGGCGCTAACTGGGATCCTCGC
W N R R D S I F Y L S K P Q T K K L R R
TGGAACAGGAGGGACAGTATTTTCTACCTTTCCAAACCGCAGACCAAGAAGCTAAGGAGA
I Y V N I L K S L L A L R P T P R S W H
ATCTATGTAAATATACTGAAATCTCTGTTGGCTCTGCGCCCAACACCCCGGAGCTGGCAC
L T L S E E R V E L D Q P L
CTCACCCTGTCTGAGGAGCGTGTAGAACTAGACCAGCCGCTATGA

**Supplementary Figure S1. ORF prediction of *circSmad1*.** A back-spliced junction-crossing ORF (peptide sequence in blue) was predicted in the 3x *circSmad1* sequence, using the NCBI ORFfinder tool (BSJ peptide highlighted in red).

**Sinha et al, Supplementary Figure S2**

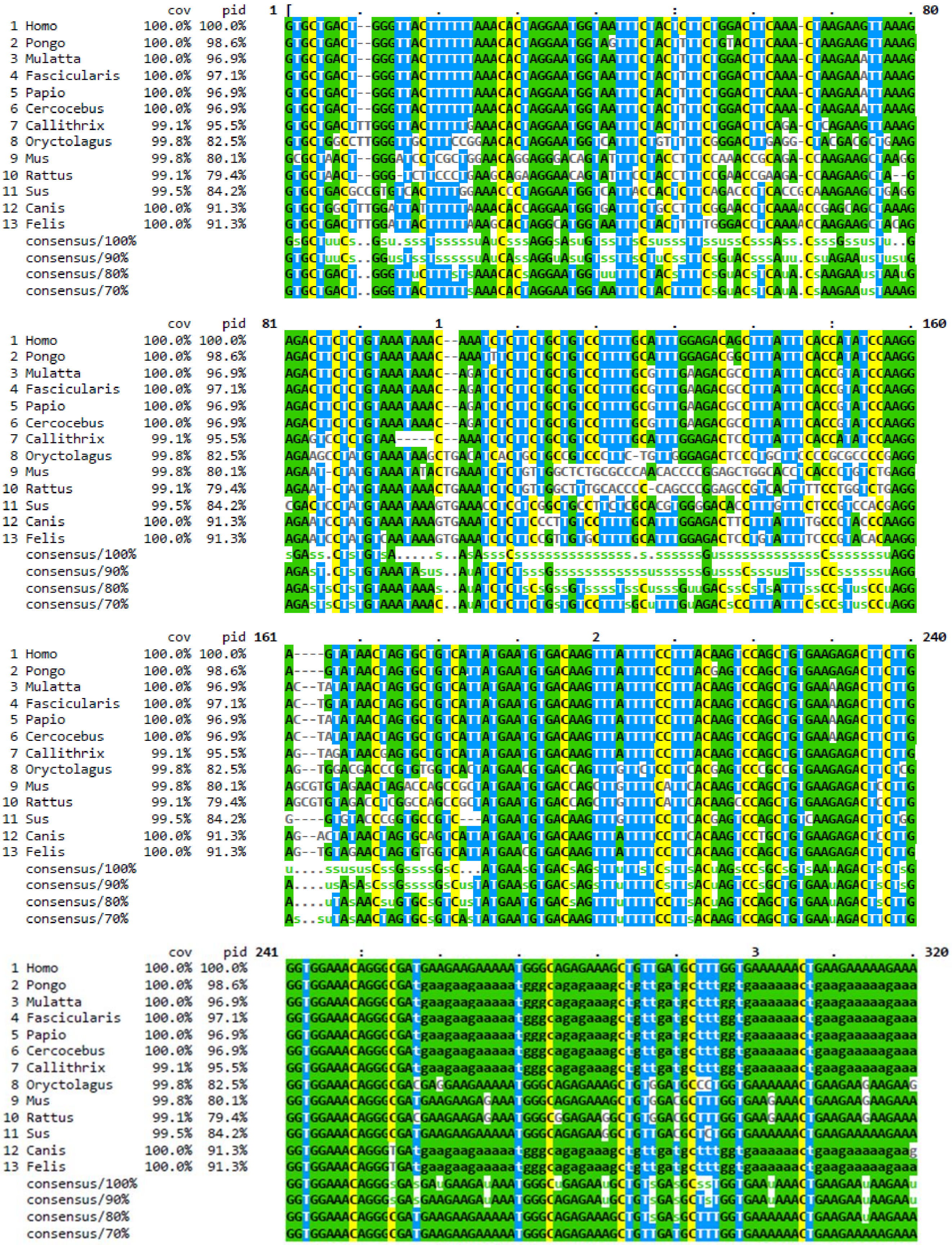

**Sinha et al, Supplementary Figure S2**

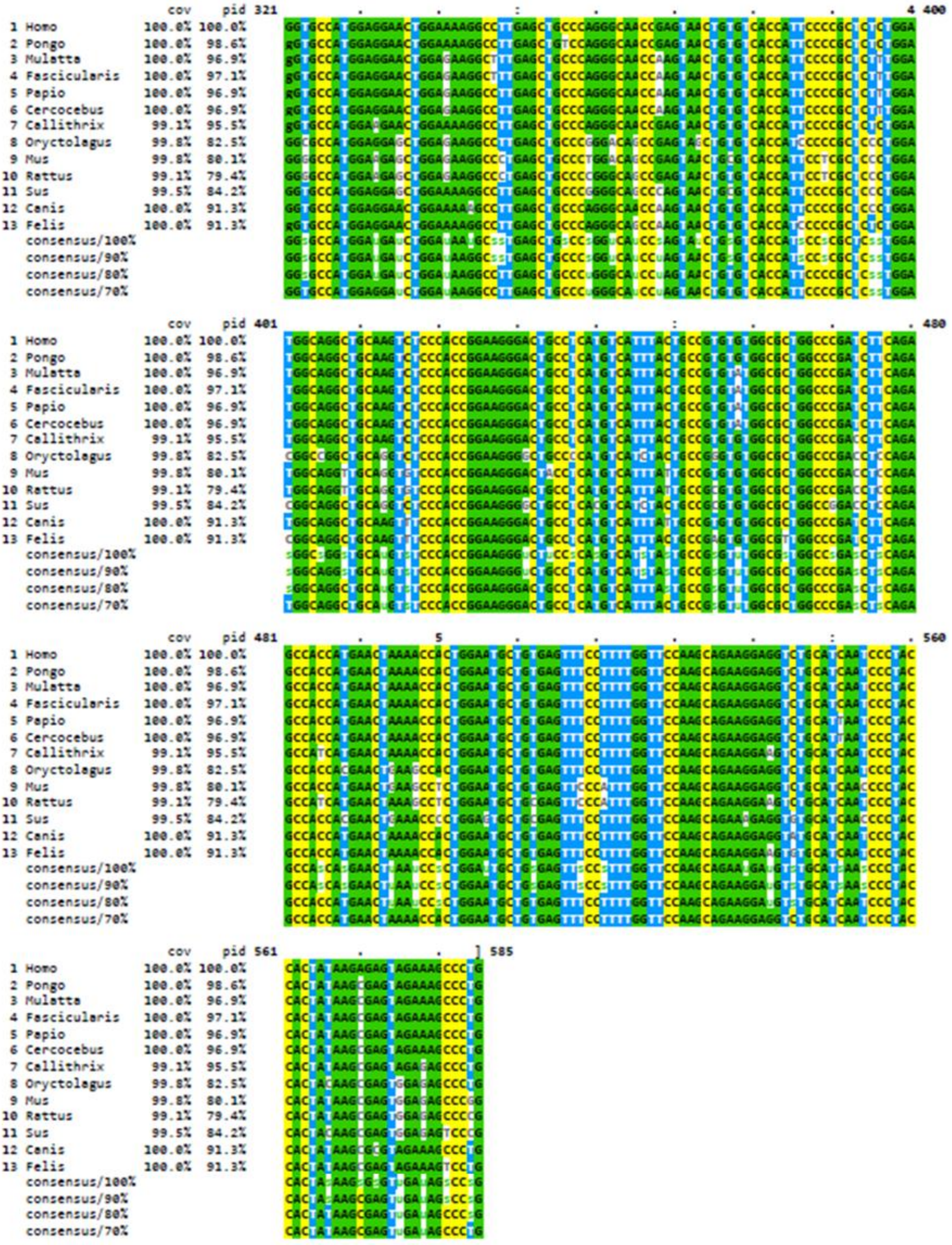


**Supplementary Figure S2. Species conservation of *circSmad1*.** Multiple sequence alignment of *circSmad1* across thirteen mammalian species curated in CIRCpedia v3 confirms evolutionary conservation of *circSmad1*.

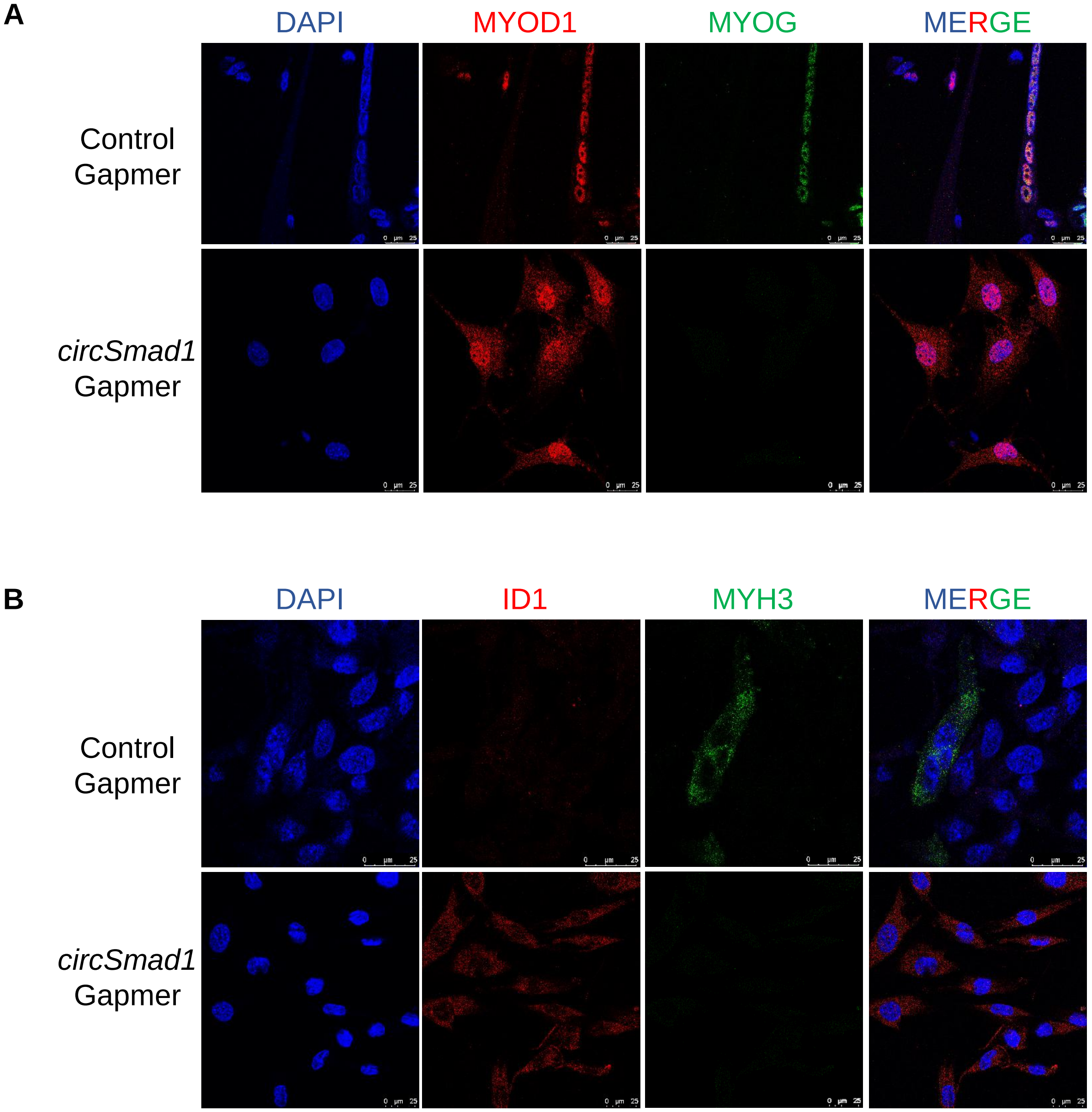


**Supplementary Figure S3. Immunofluorescence staining of MRFs in *circSmad1*-silenced C2C12 cells. A-B.** Representative confocal microscopy images showing intracellular localisation and changes in the level of MYOD1 (red), MYOG (green), ID1 (red) and MYH3 (green) in *circSmad1*-Gapmer and control Gapmer transfected C2C12 cells.

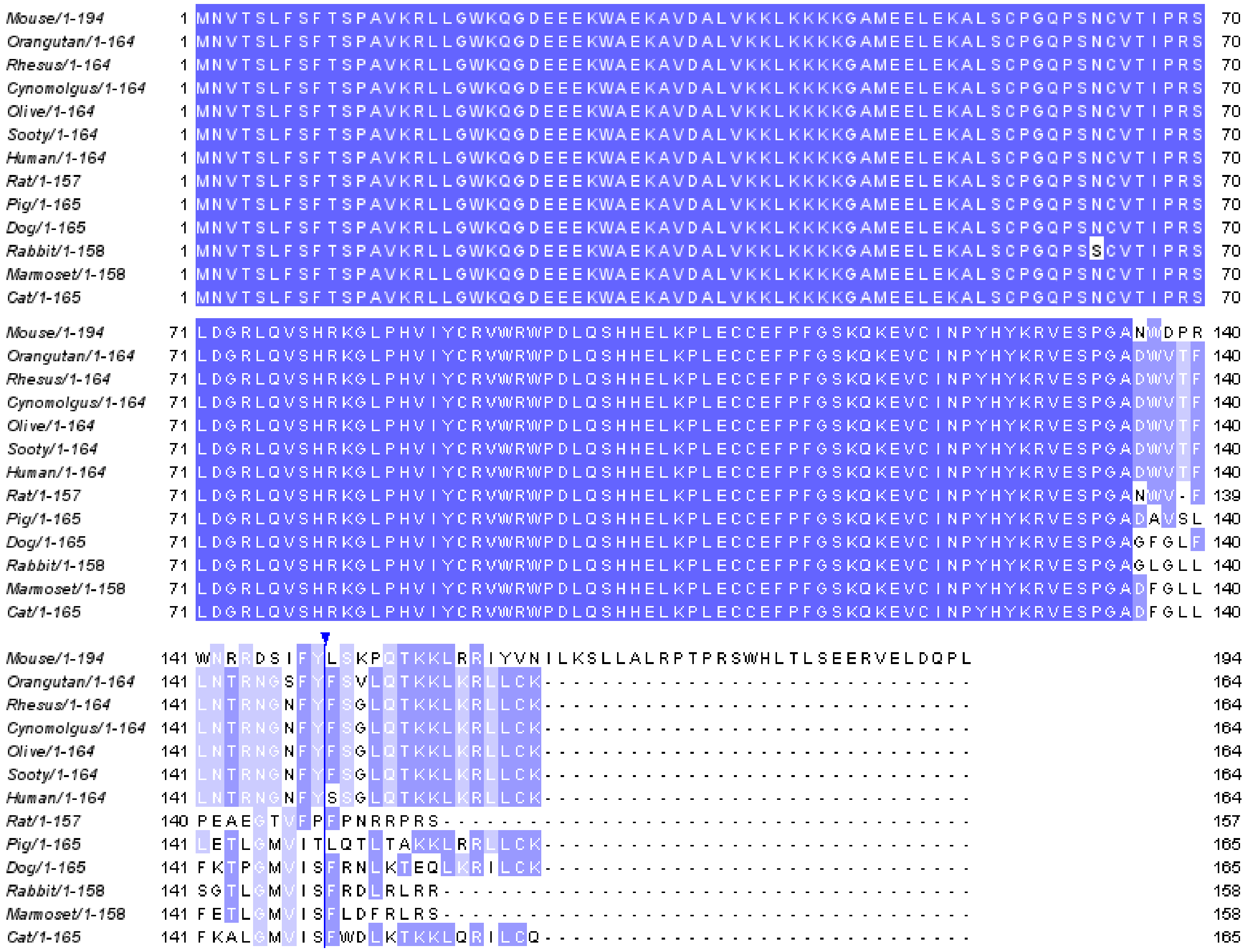

**Supplementary Figure S4. Species conservation of circSmad1-194aa.** Multiple sequence alignment of *circSmad1*-ORF across species suggests evolutionary conservation of circSmad1-194aa.

**>circSmad1-peptide**

MNVTSLFSFTSPAVKRLLGWKQGDEEEKWAEKAVDALVKKLKKKKGAMEELEKALSCPGQPSNCVTIPRSLDGRLQ
VSHRKGLPHVIYCRVWRWPDLQSHHELKPLECCEFPFGSKQKEVCINPYHYK**RVESPGANWDPR**WNRRDSIF
YLSKPQTKKLRRIYVNILKSLLALRPTPRSWHLTLSEERVELDQPL

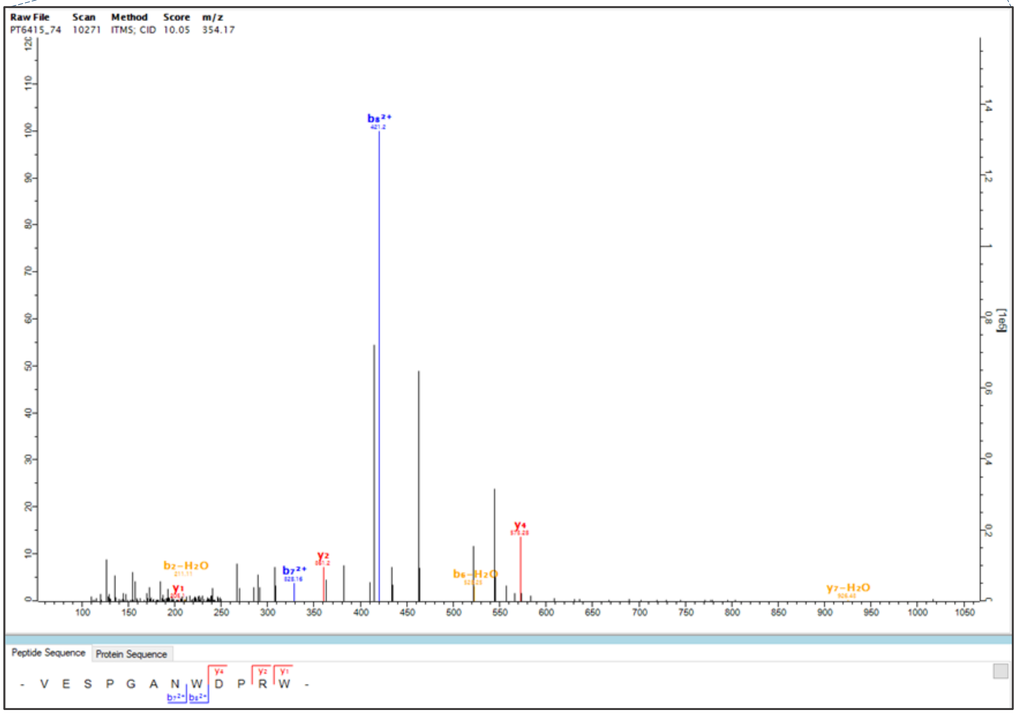


**Supplementary Figure S5. Mass spectrometry validation of circSmad1-194aa.** The junction-specific peptide of circSmad1-194aa (highlighted in pink) was detected as MS/MS spectra in mouse mass spectromentry dataset (PXD023256) using MaxQuant.

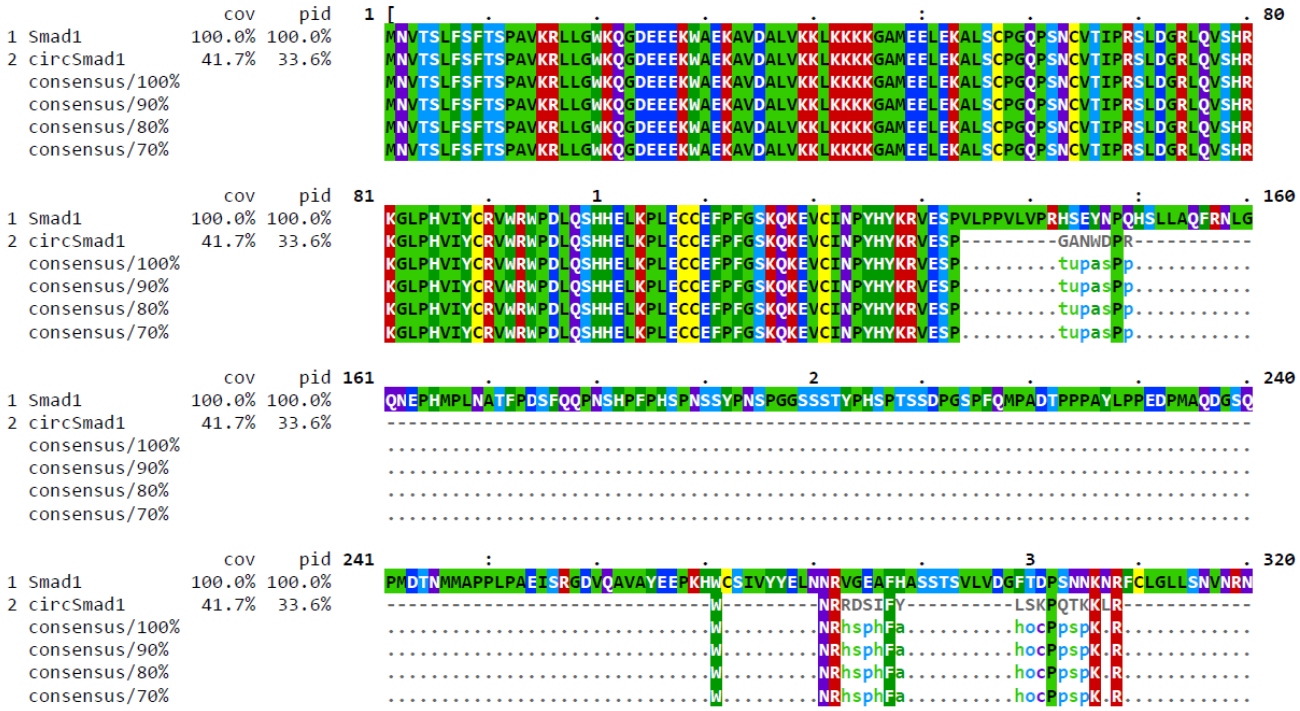


**Supplementary Figure S6. CircSmad1-194aa resembles the SMAD1-MH1 domain.** Pairwise alignment of circSmad1-194aa and host Smad1 using BLASTp showed a common MH1 domain present in circSmad1-194aa.

| Analysis | Signature accession | Signature description | Start | Stop | Score | Accession | Description | GO annotations |
|---|---|---|---|---|---|---|---|---|
| CDD | cd10490 | MH1_SMAD_1_5_9 | 9 | 132 | 6.73E-93 | - | - | - |
| SMART | SM00523 | dwAneu5 | 25 | 134 | 4.40E-68 | IPR003619 | MAD homology 1, Dwarfin-type | GO:0006355(InterPro) |
| Gene3D | G3DSA:3.90.520.10 | SMAD MH1 domain | 2 | 138 | 7.60E-59 | IPR036578 | SMAD MH1 domain superfamily | - |
| PANTHER | PTHR13703 | SMAD | 3 | 140 | 4.10E-65 | IPR013790 | Dwarfin | GO:0000978(PANTHER)\|GO:0000981(PANTHER)\|GO:0006355(InterPro)\|GO:0006357(PANTHER)\|GO:0007179(PANTHER)\|GO:0009653(PANTHER)\|GO:0030154(PANTHER)\|GO:0030509(PANTHER)\|GO:0060395(PANTHER)\|GO:0070411(PANTHER)\|GO:0071144(PANTHER) |
| FunFam | G3DSA:3.90.520.10:FF:000001 | Mothers against decapentaplegic homolog | 9 | 133 | 6.70E-79 | - | - | - |
| SUPERFAMILY | SSF56366 | SMAD MH1 domain | 13 | 131 | 1.96E-46 | IPR036578 | SMAD MH1 domain superfamily | - |
| ProSiteProfiles | PS51075 | MAD homology domain 1 (MH1) profile. | 12 | 136 | 37.080856 | IPR013019 | MAD homology, MH1 | GO:0005667(InterPro)\|GO:0006355(InterPro) |
| Coils | Coil | Coil | 34 | 54 | - | - | - | - |
| Pfam | PF03165 | MH1 domain | 31 | 131 | 1.10E-39 | IPR003619 | MAD homology 1, Dwarfin-type | GO:0006355(InterPro) |

**Supplementary Figure S7. Domain analysis of *circSmad1*-encoded peptide.** Details of functional domain sites predicted across the circSmad1-peptide sequence are described in the table, generated using the InterProScan webserver (accessed on 03-12-2025).

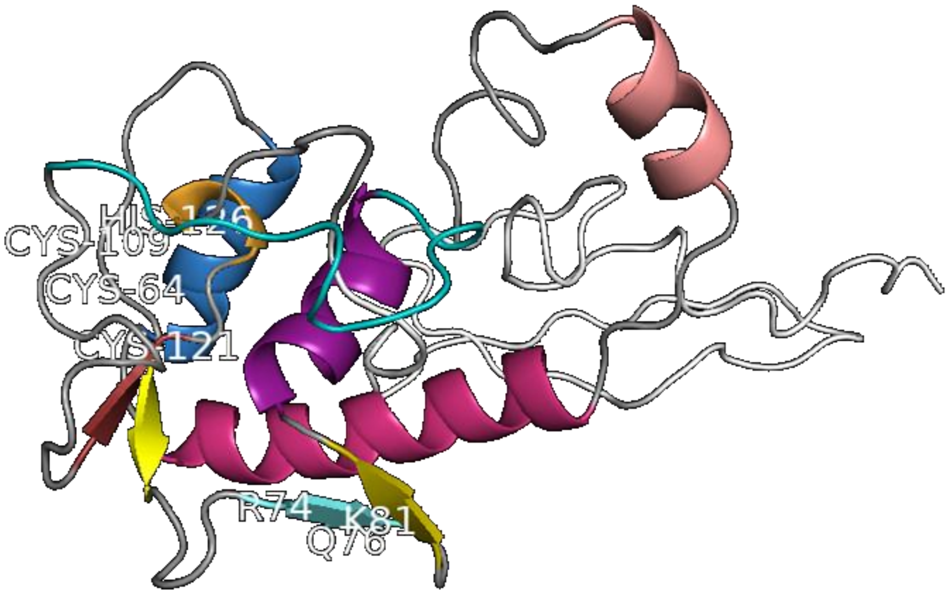


**Supplementary Figure S8. Protein structure of *circSmad1*-encoded peptide.** Three-dimensional structure of the circSmad1-peptide was predicted using AlphaFold2 Colab and visualised using PyMOL (v2.5.3) with labelling the conserved amino acid residues (Arg74, Gln76 and Lys81) present in the β-hairpin (β2–β3), which is involved in DNA binding. Another pocket of aa residues (Cys64, Cys109, Cys121 and His126), which enables zinc coordination, is also marked.

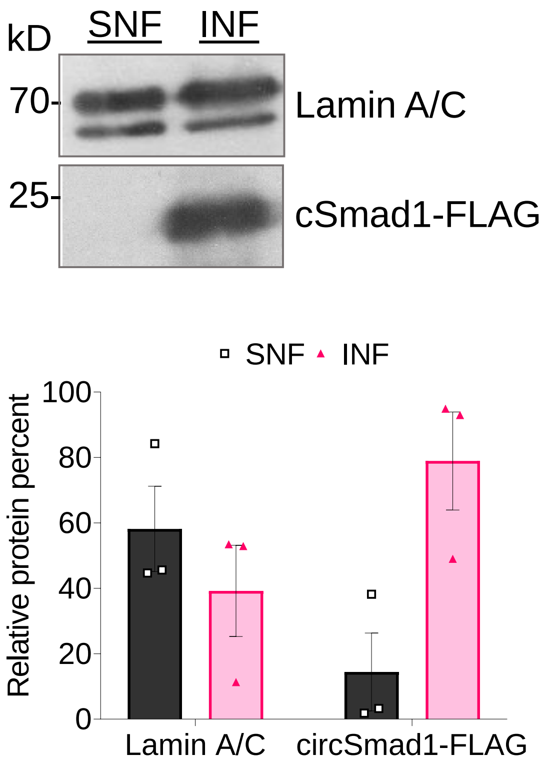


**Supplementary Figure S9. Nuclear distribution of *circSmad1*-194aa in C2C12 cells.** Representative immunoblotting image showing bands specific to Lamin A/C and circSmad1-FLAG in soluble and insoluble nuclear fractions of C2C12 myoblasts (top), and relative levels shown as bar chart, estimated by densitometry (bottom).
